# TIM3 is a context-dependent co-regulator of cytotoxic T cell function

**DOI:** 10.1101/2023.08.03.551797

**Authors:** Hanin Alamir, Carissa C.W. Wong, Amal Alsubaiti, Grace L. Edmunds, Tressan Grant, Safaa Alsulaimani, James Boyd, Christopher J. Holland, David J. Morgan, Awen M. Gallimore, Christoph Wülfing

## Abstract

TIM3 is a co-regulatory receptor that is highly expressed on multiple immune cell types, including on T cells after prolonged exposure to antigen. It marks functionally suppressed cytotoxic T lymphocytes (CTL) in the tumor microenvironment. However, it is unresolved whether TIM3 acts directly on suppressed CTL. Moreover, the nature of TIM3 ligands remains controversial. Paradoxically, TIM3 combines inhibitory function in vivo with costimulatory signaling capability in vitro. Here we have investigated TIM3 in the direct interaction of suppressed murine and human CTL with tumor target cell using spheroids. TIM3 directly inhibited the function of such CTL. TIM3 regulated the ability of suppressed CTL to polarize their cytoskeleton as a required step in cytolysis. Expression of CEACAM1 in cis, on the CTL, blocked TIM3 function, expression of CEACAM1 and galectin9 in trans, on the tumor target cells, enhanced TIM3 function. TIM3 only functioned as an inhibitory receptor on the spheroid-suppressed CTL, not on active CTL in a two-dimensional tissue culture model. These data suggest that TIM3 amplifies T cell function, serving as a co-inhibitory or co-stimulatory receptor depending on the functional context of the T cell it is expressed on.

## Introduction

CD8^+^ cytotoxic T cells (CTL) can directly kill tumor target cells. However, solid tumors often generate an immune-suppressive environment. Inhibitory receptors, CTLA-4, PD-1, TIGIT, TIM3 and LAG3, impair the anti-tumor immune response ^1, 2^. Blocking CTLA-4 and PD-1 is a cornerstone of current immunotherapy in many cancer types ^3^. However, efficacy is limited to a subset of patients and tumor types, and autoimmune side effects can be substantial. The ability of anti-PD-1 and anti-CTLA-4 to trigger the expansion of new T cell clones in draining lymph nodes is a key component of their efficacy but may also contribute to the autoimmune side effects ^4^. Therefore, therapeutic strategies that directly enhance cytolysis of tumor target cells by CTL are of interest, by themselves or as an addition to the established checkpoint blockade therapies.

Expression of the inhibitory receptor TIM3 increases with repeated T cell stimulation ^5^ reaching particularly high levels in tumors. High TIM3 expression in tumors is related to poor overall survival ^6^. Blocking TIM3 can enhance anti-tumor immunity, in particular in combination with anti-PD-1 or chemotherapy ^7^. Several basic questions in TIM3 function remain unresolved. First, does TIM3 act directly on CTL and if so how? While TIM3 is highly expressed on CD8^+^ TIL, TIM3 also regulates myeloid cell function and is highly expressed on CD4^+^ Th1 and regulatory T cells ^8, 9^. For example, blocking TIM3 on CD103^+^ dendritic cells enhances CD8^+^ T cell infiltration through increased chemokine secretion in models of breast cancer ^10^. A role of TIM3 is CD8^+^ T cell function has been established in the immune response to viral infection ^5^. In a preliminary data set, we have shown that TIM3 suppresses murine CTL killing of tumor target cells grow in three-dimensional tissue culture ^11^. However, a role of TIM3 in tumor-target cell killing by human CTL and a mechanism of action remain to be resolved. The ordered execution of a series of cytoskeletal polarization steps is critical for cytolysis and impaired in suppressed CTL ^12^. It is uncertain whether TIM3 is a regulator of such cytoskeletal polarization.

Second, functionally relevant TIM3 ligands are still controversial ^13^. TIM3 has been suggested to bind to galectin9, phosphatidylserine, high mobility group binding (HMGB) 1 and CEACAM1 both in cis and trans. HMGB1-binding sequesters DNA/HMGB1 complexes on the surface of dendritic cells to prevent activation of innate DNA sensing ^14^. Phosphatidylserine has been suggested to serve as a ligand for macrophage TIM3 in the uptake of apoptotic debris and sensitization of CTL to fratricide ^15, 16^. CEACAM1 expression in cis on a T cell enhances TIM3 function in viral infection and the anti-tumor immune response ^17^. A TIM3-Ig fusion can bind CEACAM1 expressed on Jurkat T cells ^18^, crystallographic data show a direct interaction between TIM3 and CEACAM1 N-terminal Ig domains ^17^. However, molecular binding assays fail to detect a direct interaction between TIM3 and CEACAM1 ^19^. Similarly, galectin9 has been shown to use both its N- and C-terminal carbohydrate-binding domains to trigger TIM3-dependent cell death in CD8^+^ T cells ^20^, TIM3-Ig can bind to galectin9 in an ELISA ^18^ and in surface plasmon resonance experiments using recombinant proteins ^21^. Yet, other molecular binding assays failed to detect an interaction between Galectin9 and TIM3 ^22^.

Third, while TIM3 functions as an inhibitory receptor in vivo, in vitro investigations of TIM3 signaling have consistently identified costimulatory function with increased activation of Akt, MAP kinases, Phospholipase Cγ and enhanced nuclear translocation of NFAT and NFκB ^23, 24^. The TIM3 interactome includes both stimulatory signaling intermediates, such as the p85 subunit of phosphatidylinositol 3-kinase, and inhibitory ones, such as SHP-1 and Cbl-B ^25^. It remains unresolved how inhibitory and costimulatory functions of TIM3 can be reconciled.

To address these outstanding questions with an emphasis on the role of TIM3 in CTL function, we used a recently developed murine three-dimensional tissue culture system as extended to human cells here. Murine renal carcinoma cells expressing the hemagglutinin (HA) protein from influenza virus A/PR/8 as neo-tumor-specific antigen (RencaHA) can be grown into three-dimensional spheroids. CL4 T cell receptor (TCR) transgenic CTL recognizing an HA-derived peptide can kill RencaHA target cells yet acquire a suppressed phenotype when cocultured with RencaHA spheroids. This suppressed phenotype closely resembles that of CL4 CTL isolated from RencaHA tumors in vivo ^12^. The RencaHA spheroid/CL4 CTL system thus allows the investigation of CTL suppression in the absence of other immune cells, i.e. in the direct interaction of CTL with their tumor cell targets. TIM3 inhibited tumor cell cytolysis by CTL in these spheroids. We developed a human melanoma-based spheroid system to confirm that TIM3 also suppressed the direct cytolysis of human tumor target cells by CTL. As a mechanism of TIM3 action, we found that TIM3 expression further impaired cell couple maintenance in suppressed CTL. Expression of galectin9 and CEACAM1 in the RencaHA spheroids enhanced TIM3 inhibitory function, whereas CEACAM-1 expression in the CL4 T cells blocked TIM3 function. The inhibitory function of TIM3 was restricted to interactions of CTL with tumor target cells grown as spheroids. It was lost in the interaction of CTL with the same tumor cells grown as 2D cell sheets, switching to a costimulatory function in the murine system. In spheroids CTL operated in an inhibitory signaling environment, characterized by increased expression of other inhibitory receptors on the CTL, of TIM3 ligands and by metabolic limitations. In combination our data suggest that TIM3 is a context-dependent coregulatory receptor that amplifies T cell signaling; in the inhibitory signaling environment of suppressed CTL this results in coinhibition, in active CTL this can lead to costimulation.

## Results

### TIM3 functions as a co-inhibitory receptor on CTL in tumor target cells spheroids, as a co-stimulatory receptor in 2D tissue culture

We have shown that TIM3 synergizes with adenosine in the suppression of anti-tumor immunity. Here we use the same experimental system to investigate the role of TIM3 in isolation. Renca renal carcinoma cells expressing influenza A/PR/8/H1N1 hemagglutinin (HA) induce an endogenous anti-tumor immune response and an immune-suppressive tumor microenvironment when grown subcutaneously in mice ^11, 12, 26^. The T cell receptor (TCR) of T cells from CL4 TCR transgenic mice recognizes the hemagglutinin (HA) peptide 518 to 526 (IYSTVASSL) as restricted by H2-K^d^. For simplicity and as such cells have maximal cytolytic capacity ^12^, we refer to *in vitro* agonist peptide primed CL4 CD8^+^ T cells as ‘CL4 CTL’. Upon adoptive transfer into RencaHA tumor-bearing mice, CL4 CTL infiltrate the tumor and acquire a suppressed phenotype. Incubation of *in vitro* primed CL4 CTL with RencaHA tumor cells grown as three-dimensional spheroids induces a suppressed CTL phenotype that shares key features with tumor infiltrating CL4 T cells ^12^. Extending the previous in vivo tumor growth data, two groups of BALB/c mice bearing 12-day old RencaHA tumors were treated with the anti-TIM3 blocking mAb RMT3-23. To provide a standardized number of anti-tumor CTL at a comparable stage of tumor growth, we used two i.v. adoptive T cell transfers of 5×10^6^ CL4 TCR transgenic CTL on days 12 and 14. Tumor growth at the time of adoptive transfer and during tumor regrowth two to three weeks later was reduced upon treatment with anti-TIM3 (Fig. 1A), confirming inhibitory TIM3 function in vivo in our murine experimental system.

**Figure 1.**
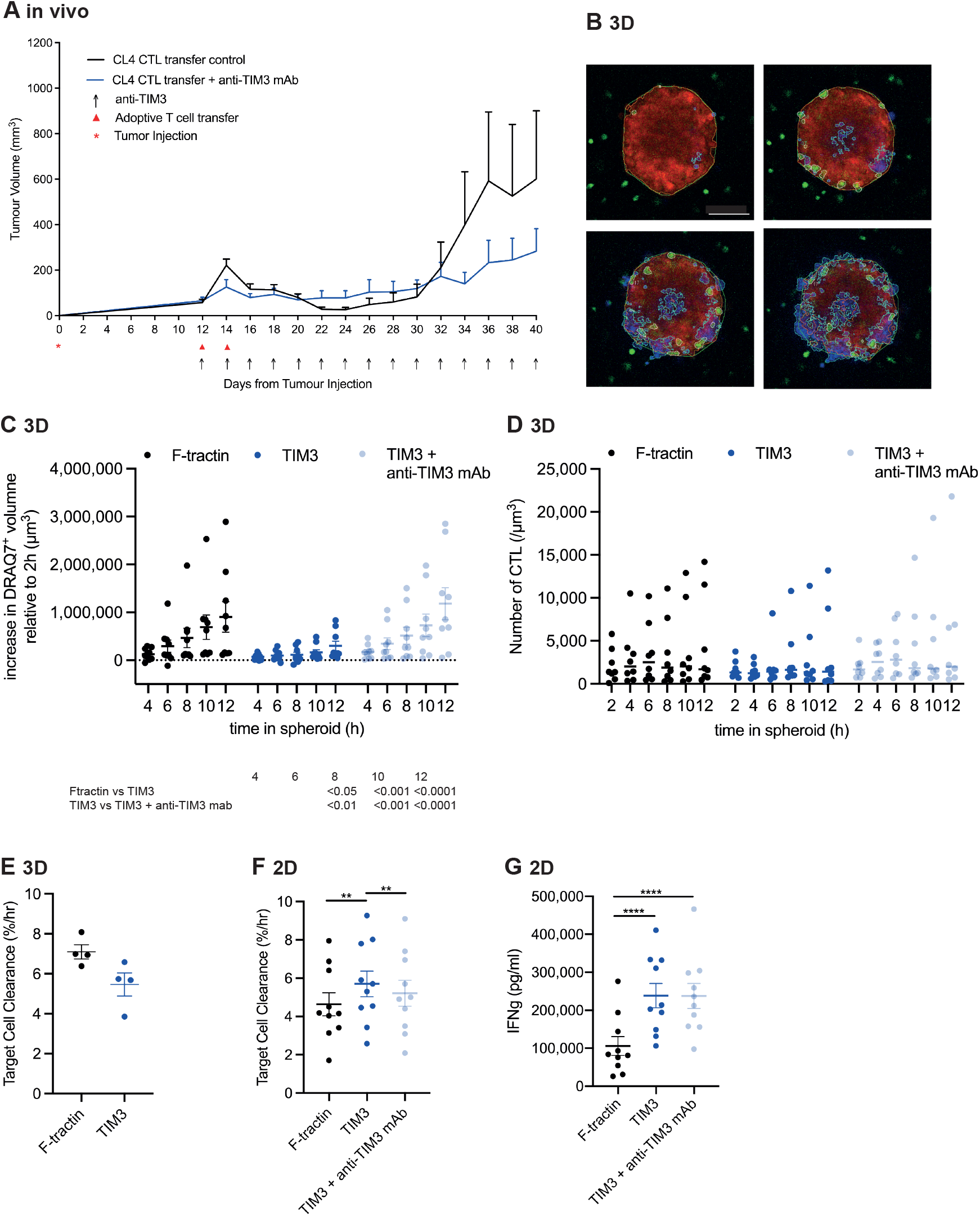
TIM3 suppresses CTL function in a three-dimensional tissue context. **A** RencaHA tumor-bearing BALB/c mice were injected i.v. twice with 5 x10^6^ purified CL4 CTL on days 12 and 14 and treated with anti-TIM3 mAb (clone RMT3-23) or isotype-control antibody, as shown. Tumor growth is displayed as mean tumor volumes + SEM with N=13 mice over 2 independent experiments, with at least 6 mice per group. **B** Representative images of a RencaHA tdTomato spheroid (red) with F-tractin-GFP-expressing CL4 CTL (green) as stained for cell death with DRAQ7 (blue) over time of CTL spheroid interaction. Segmentations for volume measurements are shown. scale bar=100µm. **C, D** CL4 CTL retrovirally transduced to express TIM3-GFP or F-tractin-GFP cocultured with RencaHA tdTomato spheroids incubated with K^d^HA peptide for 12h ± 10µg/ml anti-TIM3 mAb (clone RMT3-23) with images acquired every 2h. Each data point is an independent experiment (N=9) with a total of 47-50 spheroids analyzed per condition. C Spheroid death, as measured by the increase in DRAQ7^+^ spheroid volume, is shown. Significance of differences between conditions at indicated time points is given in the table below. Single spheroid data are given in Fig. S1G. D SIL densities are shown with the mean ± SEM for the same experiments as in C. No significant differences. **E** *In vitro* killing of KdHA-pulsed Renca tumor cells by CL4 SIL expressing TIM3-GFP or F-tractin-GFP. Average cell death rates calculated as percentage decrease in area covered by tumor cells per hour are means ± SEM from 4 independent experiments. **F** *In vitro* killing of KdHA-pulsed Renca tumor cells by CL4 CTL expressing F-tractin-GFP, TIM3-GFP or TIM3-GFP in the presence of anti-TIM3 mAb. Average cell death rate calculated as percentage decrease in area covered by tumor cells per hour are means ± SEM from 10 independent experiments. **G** IFNγ amounts in the supernatants of the killing assays in F at the end of the assay. ** p < 0.01, **** p < 0.0001; p values calculated using one-way (E-G) and two-way (C, D) ANOVA

To determine whether TIM3 directly regulates the interaction of suppressed CTL with their tumor target cells, we employed the *in vitro* reconstruction of CL4 CTL suppression in Renca spheroids (Fig. 1B). Spheroids were grown to a diameter of around 250µm, had a necrotic core but were not hypoxic (Fig. S1A-C). Tumor cell killing in spheroids was dependent on the presence of the CL4 TCR (Fig. S1D). The majority of spheroids-associated CTL were firmly bound to the spheroid surface or penetrated slightly, less than 8µm deep (Fig. S1E). Accordingly, Renca cell death was greatest at the spheroid periphery (Fig. S1F). To determine roles of TIM3, we expressed TIM3-GFP in CL4 CTL using cell sorting to adjust expression to 5×10^5^ molecules per T cell ^27^. As control we expressed the F-actin-binding peptide F-tractin ^28^ linked to GFP at the same level. TIM3 is increasingly upregulated upon repeated and persistent stimulation of human and murine T cells with antigen, reaching levels of more than 100-fold above unstimulated ^5, 29, 30, 31^. In the sorted TIM3-GFP expressing CL4 CTL TIM3 expression was elevated 150-fold (Fig. S2A-C), thus representing TIM3 expression on extensively stimulated CTL.

As a measure of CL4 CTL killing in Renca spheroids, the increase in the volume of dead tumor cells in Renca spheroids between 2h and 12h of incubation with CL4 CTL was reduced from 910,000±320,000µm^3^ upon F-tractin expression to 300,000±100,000µm^3^ upon TIM3 expression (p<0.0001) (Figs. 1C, S1G). Establishing specificity, tumor cell killing was restored in the presence of the TIM3 blocking mAb RMT3-23 to an increase in the volume of dead tumor cells at 12h of 1,200,000±340,00µm^3^ (Fig. 1C). CTL infiltration into the tumor spheroids did not differ (Fig. 1D), suggesting that the execution of cytolysis was inhibited by TIM3 at the single CTL level. To directly test this suggestion, we isolated CTL after spheroid incubation and determined killing of the same Renca tumor target cells incubated with a HA agonist peptide concentration of 2µg/ml in an overnight imaging-based killing assay ^12^. Tumor cell killing was reduced upon TIM3 expression by 25% (Fig. 1E). As some function of suppressed CTL is recovered after removal from a suppressive environment ^32^, the smaller effect size relative to the in-spheroid killing is to be expected. Together these data establish that TIM3 suppresses CTL killing of tumor cells in spheroids.

In a further simplification of the physiological context, we determined whether TIM3 inhibited tumor target cell killing in the interaction of CL4 CTL with a two-dimensional sheet of the same Renca target cells. In contrast to the spheroid data, expression of TIM3 increased killing of Renca target cells from 4.6±0.6%/h reduction in the plate bottom area covered by tumor cells as our imaging-based measure of killing to 5.7±0.7%/h (p<0.01) (Fig. 1F). The enhancement of killing upon TIM3 expression was partially reversed by the anti-TIM3 mAb RMT3-23 (5.2±0.7%/h area reduction, p<0.005 versus TIM3 expression) (Fig. 1F). Co-stimulation of CTL function upon TIM3 expression extended to IFNγ secretion as a second key effector function of CTL. In killing assay supernatants at the end of the killing assay, TIM3 expression more than doubled IFNγ amounts from 105±25ng/ml to 240±30ng/ml (p<0.0001) (Fig. 1G) as not reversed by the TIM3-blocking mAb (Fig. 1G). Thus, using the same CTL and tumor target cells, TIM3 switched from co-inhibitory to co-stimulatory function when tumor cells were grown in 3D versus 2D tissue culture.

To confirm a context-dependent role of TIM3 across experimental systems and species, we expressed TIM3-GFP together with the 1G4 TCR by lentiviral transduction in primary human CD8^+^ T cells (Fig. S2A-C). The 1G4 TCR recognizes the melanoma-derived NY-ESO-1 peptide 157 to 165 (SLLMWITQC) as restricted by HLA-A2*02:01. TIM3 was already expressed upon human CD8^+^ T cell culture with anti-CD3/CD28 beads and IL-2 as described before ^29^. Upon lentiviral transduction of these cells with TIM3-GFP, TIM3 expression was further elevated 2.4-fold reaching a level of 40-fold above the TIM3-negative portion of the human and murine CTL cultures (Fig. S2B, C). This was the highest level of human TIM3 expression that we could technically achieve, representing TIM3 levels on stimulated CTL but below the maximum obtained with repeated persistent stimulation and TIM3-GFP expression in the CL4 CTL. In the interaction of the 1G4 CTL with spheroids of the melanoma cell line Mel624, the increase in the volume of dead tumor cells between 2h and 12h of CTL incubation was reduced from 130,000±8,000µm^3^ upon expression of F-tractin-GFP as a control to 20,000±5,000 µm^3^ (p<0.01) upon expression of TIM3-GFP (Figs. 2A, B, S2D), confirming a strong inhibitory role of TIM3. In contrast, in the interaction of the same human 1G4 CTL with the same Mel624 tumor cells cultured as a 2D cell sheet and incubated with 2µg/ml of the NY-ESO-1 peptide, tumor cell killing and IFNγ secretion were unaffected by expression of TIM3, as confirmed with Hep-G2 hepatocellular carcinoma target cells (Fig. 2C, D). Thus, TIM3 was only inhibitory in the 3D spheroid context, as in the mouse experiments. The lack of a complete switch to a costimulatory effect in 2D may be related to the lower human TIM3 expression level.

**Figure 2.**
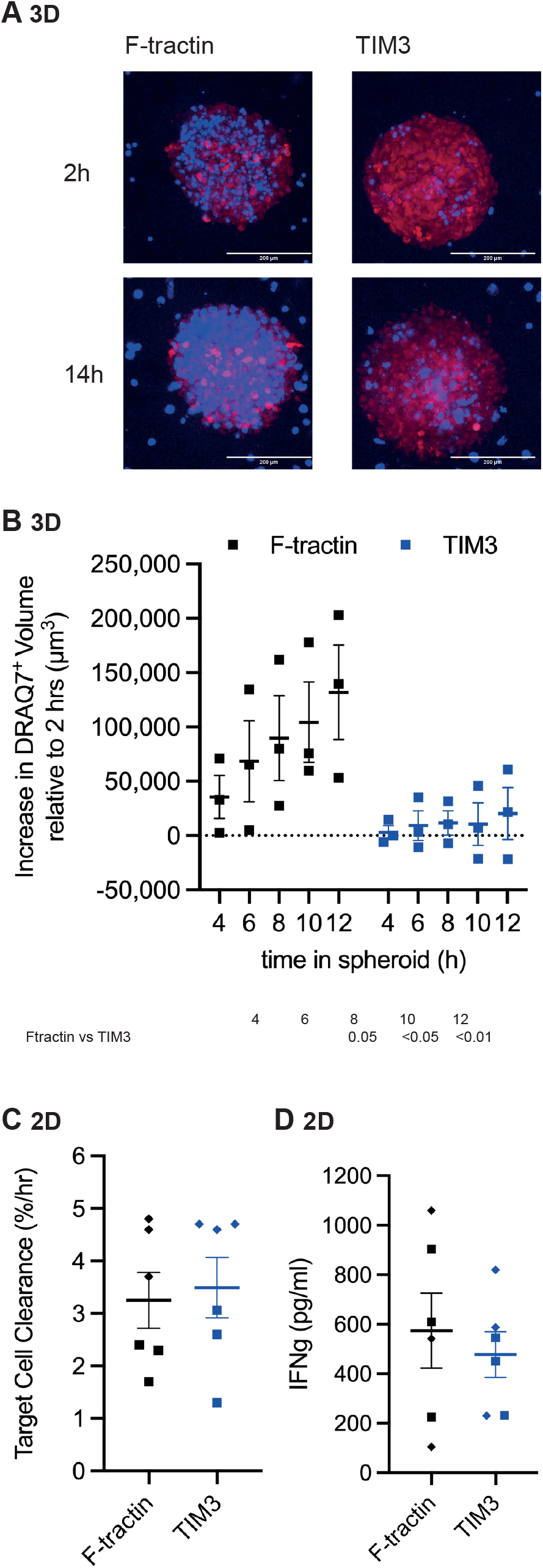
TIM3 suppresses human CTL function in a three-dimensional tissue context. **A** Representative images of Mel624 tdTomato spheroids (red) as stained for cell death with DRAQ7 (blue) over time of spheroid interaction with human CTL expressing the 1G4 TCR together with F-tractin-GPF or TIM3-GFP as indicated. **B** Human CTL lentivirally transduced to express the 1G4 TCR together with TIM3-GFP or F-tractin-GFP cocultured with Mel624 tdTomato spheroids incubated with NY-ESO-1 peptide for 12h with images acquired every 2h. Each data point is an independent experiment (N=3) with a total of 14 spheroids analyzed per condition. Spheroid death, as measured by the increase in DRAQ7^+^ spheroid volume, is shown. Significance of differences between conditions at indicated time points is given in the table below as calculated using 2-way ANOVA. Single spheroid data are given in Fig. S2D. **C** *In vitro* killing of NY-ESO-1-pulsed Mel624 (square symbols) or Hep-G2 (rhomboid symbols) tumor cells by 1G4 CTL expressing TIM3-GFP or F-tractin-GFP. Average cell death rates calculated as percentage decrease in area covered by tumor cells per hour are means ± SEM from 3 independent experiments for each tumor cell line. **D** IFNγ amounts in the supernatants of the killing assays in C at the end of the assay. p values calculated using two-way ANOVA

### TIM3 regulates the maintenance of cell couples between CTL and tumor target cells

The inability to effectively execute a series of cytoskeletal polarization steps required for effective cytolysis characterizes suppressed CTL ^12^. To determine such polarization, we imaged the interaction of CL4 spheroid-infiltrating lymphocytes (CL4 SIL) expressing TIM3 with Renca tumor target cells in comparison to that of *in vitro* CL4 CTL. The formation of a tight cell couple upon initial contact of a CTL with a tumor target cell is the first step of cytolysis (Fig. 3A). CL4 SIL cell coupling was inefficient (35±0.5%) and further slightly reduced upon TIM3 expression (29±1%). In contrast, TIM3 expression in CL4 CTL enhanced cell coupling from 64±7% to 78±4% (p=0.001)(Fig. 3B). One hallmark of defective cell couple maintenance of suppressed CTL are lamellae directed away from the cellular interface (Fig. 3A). The frequency of CL4 SIL with off-interface lamellae was high (91±5%) as previously established ^12^ and thus could not be further enhanced upon TIM3 expression (Fig. 3C). In contrast, TIM3 expression in CL4 CTL significantly reduced the frequency of T cells with such lamellae from 70±4% to 46±10% (p=0.05) (Fig. 3C). The time to first occurrence of an off-interface lamella as another measure of cell couple instability was short in SIL as previously established ^12^, slightly further reduced upon TIM3 expression in CL4 SIL, longer in CL4 CTL and delayed upon TIM3 expression (Fig. 3D). Another hallmark of suppressed CTL is T cell translocation over the tumor cell surface away from the site of initial coupling (Fig. 3A). In CL4 SIL expressing TIM3, 44±7% of cell couples showed such translocation in contrast to only 7±5% of CL4 SIL expressing F-tractin (p=0.001))(Fig. 3E). Such translocation was virtually absent in CL4 CTL (1.5±1.5%)(Fig. 3E). Together these data establish that TIM3 expression destabilized cell couples in CL4 SIL while stabilizing them in CL4 CTL, consistent with a co-inhibitory versus co-stimulatory role in CL4 SIL versus CL4 CTL, respectively. TIM3 thus consistently regulated cell couple maintenance in the interaction of CTL with tumor target cells as an element of its mechanism of action.

**Figure 3.**
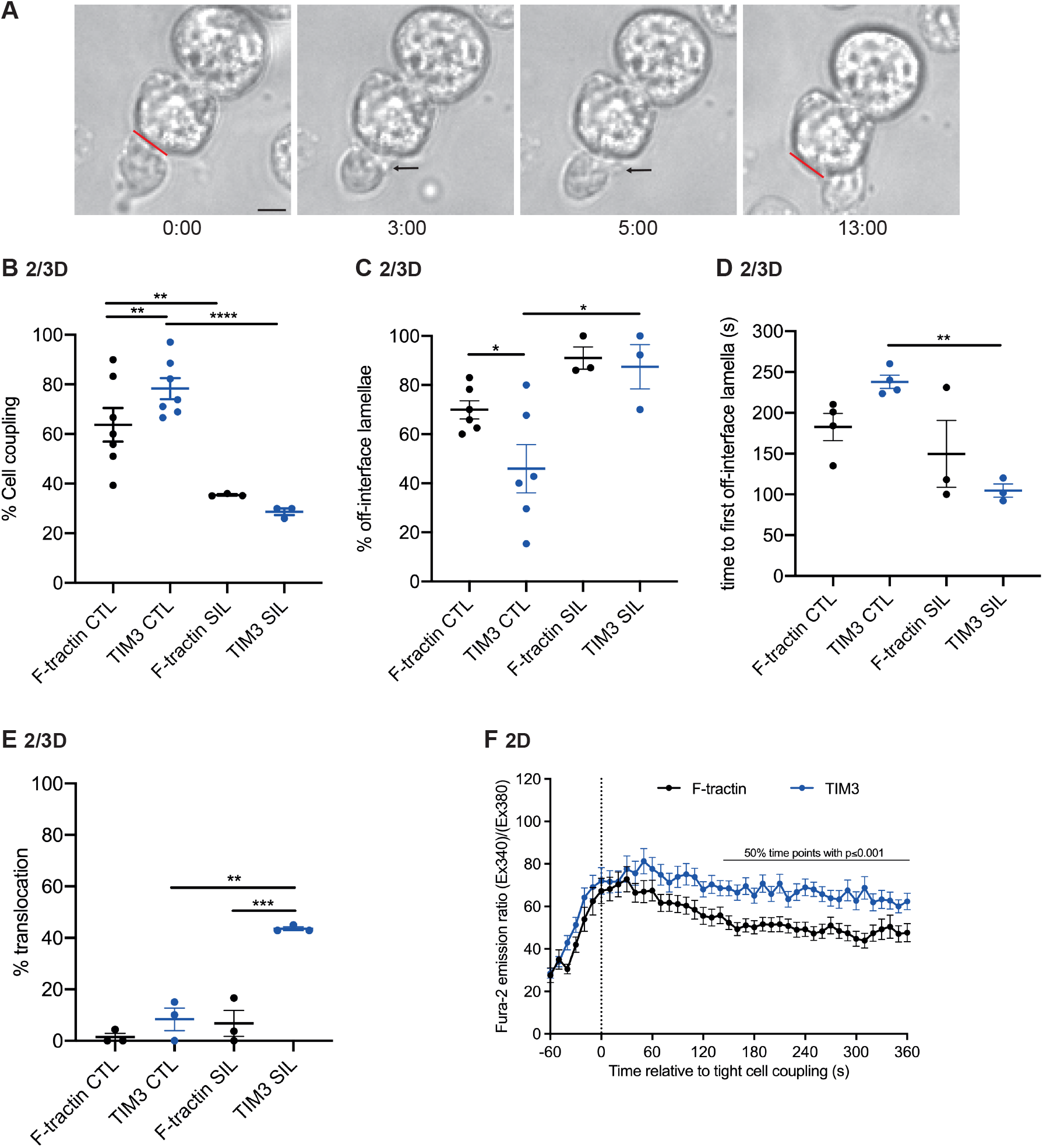
SIL cell couple maintenance is impaired as regulated by TIM3. **A-E** Microscopy analysis of interactions over time of Clone 4 CTL and SIL with HA peptide pulsed Renca cells. A Images of a CL4 SIL expressing TIM3-GFP are representative of ≥3 independent experiments. Red line indicates original position of tight-cell couple formation on Renca cell. Arrows indicate off-synapse lamellae. Time in min relative to tight cell couple formation. Scale bar=5µm. B-E Percentage of Clone 4 CTL and SIL expressing F-tractin-GFP or TIM3-GFP that form a tight cell couple upon contact with a tumor target cell (B) with off-interface lamellae (C), time of first off-synapse lamella (D) and translocation of more than one immune synapse diameter (E). Data are means ± SEM from ≥ 3 independent experiments. Cell couples were imaged for 10±0.5min (at least 2min). **F** CL4 CTL expressing F-tractin-GFP or TIM3-GFP interacted with Renca cells incubated with 2µg/ml K^d^HA peptide. The ratio of Fura-2 emissions at 510nm upon excitation at 340nm over 380nm is given relative to time of tight cell coupling as the mean ± SEM. N=5 independent experiments, 54, 68 T cells analyzed per condition. Single cell and individual experiment average data are given in Fig. S3. * p < 0.05, ** p < 0.01, *** p<0.001, **** p < 0.0001; p values calculated using proportions z-test (B, C, E), one-way ANOVA (D) and student’s t-test (F for each time point)

An important proximal signaling step in T cell activation is the elevation of the cytoplasmic calcium concentration. It was further elevated upon TIM3 expression in CL4 CTL later than 150s after tight cell coupling when stimulated with 2µg/ml HA peptide at the border of the stringent Bonferroni-corrected p-value of 0.001 (Figs. 3F, S3), consistent with a costimulatory function of TIM3 in the 2D context. Sufficient numbers of TIM3-expressing CL4 SIL for an analogous SIL experiment could not be obtained.

### TIM3 co-regulatory function is largely dependent on its cytoplasmic domain

To determine a role of TIM3-induced signal transduction in TIM3 function, we expressed a TIM3 variant lacking the cytoplasmic domain in CL4 CTL at the same level as full-length TIM3. CL4 CTL-mediated killing of tumor target cells in tumor cell spheroids was doubled upon deletion of the TIM3 cytoplasmic domain, an almost complete reversion of the TIM3 inhibitory effect to levels seen upon anti-TIM3 mAb treatment (Figs. 4A, S4A). CL4 CTL infiltration into spheroids was not substantially changed (Fig. S4B). The enhancement of IFNγ secretion in 2D tissue culture by TIM3 was also reversed upon deletion of the TIM3 cytoplasmic domain, almost completely at limiting agonist peptide concentrations (Fig. 4B). The enhancement in cytolytic killing was only partially reversed (Fig. 4C). Together these data suggest that signals induced by the TIM3 cytoplasmic domain play a substantial role in TIM3 co-regulation of most elements of CTL effector function.

**Figure 4.**
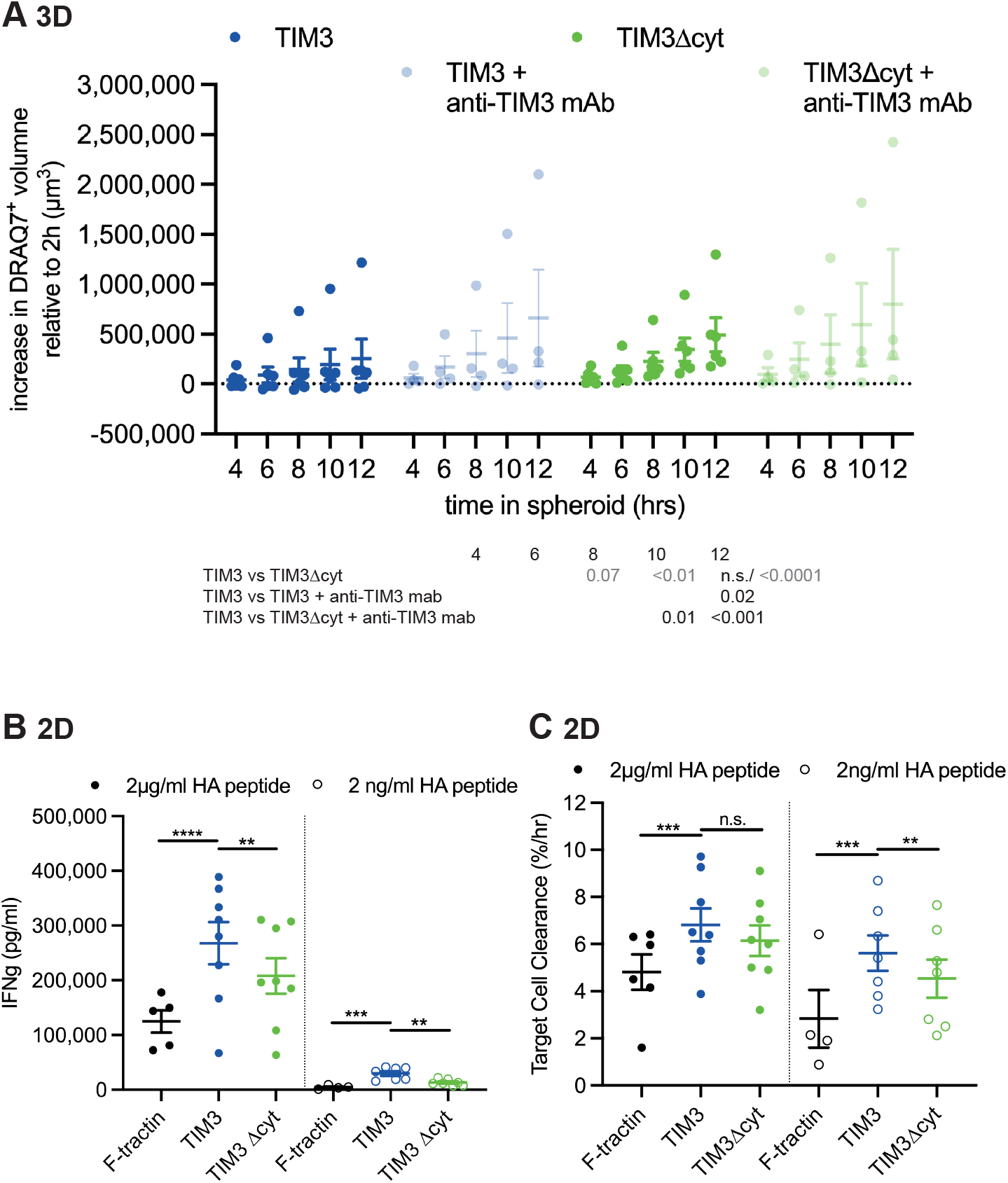
TIM3 co-regulatory function is only partially dependent on its cytoplasmic domain. **A** CL4 CTL retrovirally transduced to express TIM3-GFP or TIM3Δcyt-GFP cocultured with RencaHA tdTomato spheroids incubated with K^d^HA peptide for 12h ± 10µg/ml anti-TIM3 mAb (clone RMT3-23) with images acquired every 2h. Each data point is an independent experiment (N=4-6) with a total of 11-23 spheroids analyzed per condition. Spheroid death, as measured by the increase in DRAQ7^+^ spheroid volume, is shown. Significance of differences between conditions at indicated time points is given in the table below, mixed effect analysis of the entire data in black, two-way ANOVA of the comparison TIM3-GFP versus TIM3Δcyt-GFP in grey. Single spheroid and spheroid infiltration data are given in Fig. S4. **B, C** *In vitro* killing of Renca tumor cells pulsed with 2µg/ml or 2ng/ml KdHA peptide by CL4 CTL expressing F-tractin-GFP, TIM3-GFP or TIM3Δcyt-GFP. C Average cell death rate calculated as percentage decrease in area covered by tumor cells per hour are means ± SEM from 4-8 independent experiments. B IFNγ amounts in the supernatants of the killing assays in C at the end of the assay. ** p < 0.01, *** p<0.001; p values calculated using one-way ANOVA (B, C, separately for each agonist peptide concentration) and mixed effect analysis (A)

### CEACAM1 expression in cis inhibits TIM3 function

CEACAM1 is highly expressed alongside TIM3 in exhausted T cells ^17^. To investigate how CEACAM1 expression on CTL regulates TIM3 function, we used retroviral transduction and P2A ribosome hopping to express TIM3-GFP and CEACAM1 in parallel in CL4 CTL (Fig. 5A). Based on GFP fluorescence in the CL4 CTL sorted for TIM3-GFP, TIM3-GFP expression in these cells was 2-fold lower than in CL4 T cells expressing TIM3-GFP only. However, cell surface expression of TIM3 was reduced 9.5-fold upon co-expression of CEACAM1 (Fig. S5A, B), suggesting that CEACAM1 expression in cis impairs TIM3 trafficking to the plasma membrane and/or enhances its internalization from the cell surface. As a control we also generated CL4 CTL expressing CEACAM1-GFP only. Expression of CEACAM1 in isolation did not change tumor target cell death in Renca spheroids (Figs. 5B, S5C). However, expression of CEACAM1 alongside TIM3 largely reversed the inhibitory effect of TIM3 only expression (Fig. 5B). CL4 T cell infiltration into spheroids was unchanged (Fig. S5D). These data are consistent with two scenarios, a loss of TIM3 function by reduced cell surface expression and/or blockade of TIM3 ligand binding by CEACAM1.

**Figure 5.**
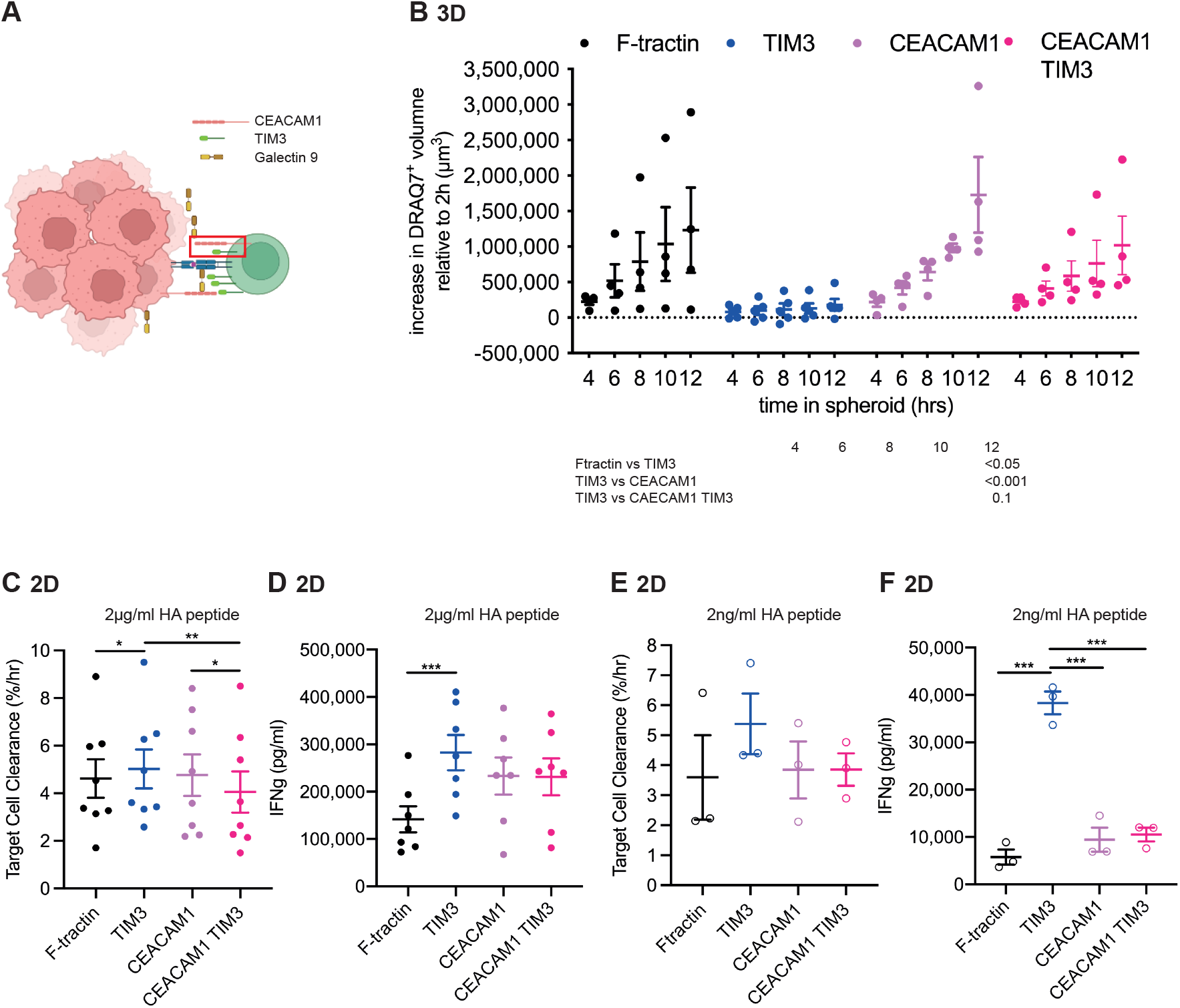
CEACAM1 expression in cis inhibits TIM3 function. **A** Scheme of the interaction of CTL with spheroids with TIM3 and potential ligands labelled. Red box shows an interaction between TIM3 and CEACAM1 on CTL in cis. Figure created with BioRender.com. **B** CL4 CTL retrovirally transduced to express F-tractin-GFP, TIM3-GFP, CEACAM1-GFP or TIM3-GFP together with CEACAM1 cocultured with RencaHA tdTomato spheroids incubated with K^d^HA peptide for 12h with images acquired every 2h. Each data point is an independent experiment (N=4) with a total of 12-15 spheroids analyzed per condition. Spheroid death, as measured by the increase in DRAQ7^+^ spheroid volume, is shown. Significance of differences between conditions at indicated time points is given in the table below. Single spheroid and spheroid infiltration data are given in Fig. S5C, D. **C-F** *In vitro* killing of Renca tumor cells pulsed with 2µg/ml or 2ng/ml KdHA peptide by CL4 CTL expressing F-tractin-GFP, TIM3-GFP, CEACAM1-GFP or TIM3-GFP together with CEACAM1. C, E Average cell death rates calculated as percentage decrease in area covered by tumor cells per hour are means ± SEM from 3-8 independent experiments. D, F IFNγ amounts in the supernatants of the killing assays in C, E at the end of the assay. * p<0.05, ** p < 0.01, *** p<0.001, p values calculated using one-way (C-F) and two-way (B) ANOVA

We also investigated the CL4 CTL interaction with Renca tumor target cells in 2D tissue culture. The co-stimulatory effect of TIM3 expression on target cell killing (Fig. 1E) was reversed by CEACAM1 co-expression (Fig. 5C), as was the effect on IFNγ secretion, albeit only partially (Fig. 5D). The size of the effect of TIM3 expression on CL4 CTL function becomes larger when the HA agonist peptide concentration is reduced to 2 ng/ml (Figs. 5E, F), matching the MHC/peptide concentration achieved by endogenous HA processing (Fig. S5E). While upon stimulation with 2ng/ml HA peptide TIM3 expression in CL4 CTL increased IFNγ secretion from 6±1.5 to 33±2.5 ng/ml (p<0.001), co-expression of CEACAM1 with TIM3 on the CL4 CTL only yielded 10.5±1.5 ng/ml IFNγ, not significantly different from the F-tractin control (Fig. 5F). The 2D data thus corroborate that CEACAM1 expression on CL4 CTL in cis interferes with TIM3 function.

### CEACAM1 in trans enhances TIM3 function

CEACAM1 can be expressed by tumor and immune cells and could serve as a ligand for TIM3 in trans ^33^ (Fig. 6A). To investigate how CEACAM1 expression in trans regulates TIM3 function, we transfected Renca cells to express CEACAM1-tdTomato. While Renca cells did not express CEACAM-1 endogenously, neither in tissue culture not when isolated from subcutaneous Renca tumors (Fig. S6A, B), the CEACAM-1 expression on CEACAM1-tdTomato transfected Renca cells was only 51% above CEACAM-1 expression on CD45^+^ tumor cells (Fig. S6A, B). Such overexpression of CEACAM1 on the Renca cells changed spheroid morphology from round to more elongated (Fig. 6B). These data suggest a change in tumor cell biology upon CEACAM1 overexpression, as not further explored within our emphasis on CTL function here. While the resulting increase in the surface area to volume ratio of the Renca CEACAM1 spheroids should facilitate cytolysis through improved CTL access, tumor cell killing was moderately inhibited in Renca CEACAM1 spheroids interacting with CL4 CTL expressing TIM3 (Figs. 6C-E, S6C). This inhibition could be reversed with a blocking antibody against TIM3 (Fig. 6D). Spheroid infiltration was unaltered (Fig. S6D). These data establish that CEACAM1 at near physiological high expression levels in trans serves as an inhibitory ligand for TIM3 in the three-dimensional spheroid context. When CL4 CTL interacted with Renca target cells in 2D, CEACAM1 overexpression on the Renca cells yielded a moderate increase in IFNγ generation (Fig. 6F), consistent with CEACAM1 serving as a costimulatory ligand for TIM3. However, target cell killing was significantly (p<0.001) impaired upon CEACAM1 overexpression (Fig. 6G). This impairment could not be reversed with antibodies blocking TIM3 (Fig. 6G), suggesting that it was not a role of CEACAM1 as a ligand for TIM3 that inhibited target cell killing. Changes in tumor cell biology are conceivable. For example, CEACAM1 is an established regulator of tumor cell migration ^34^.

**Figure 6.**
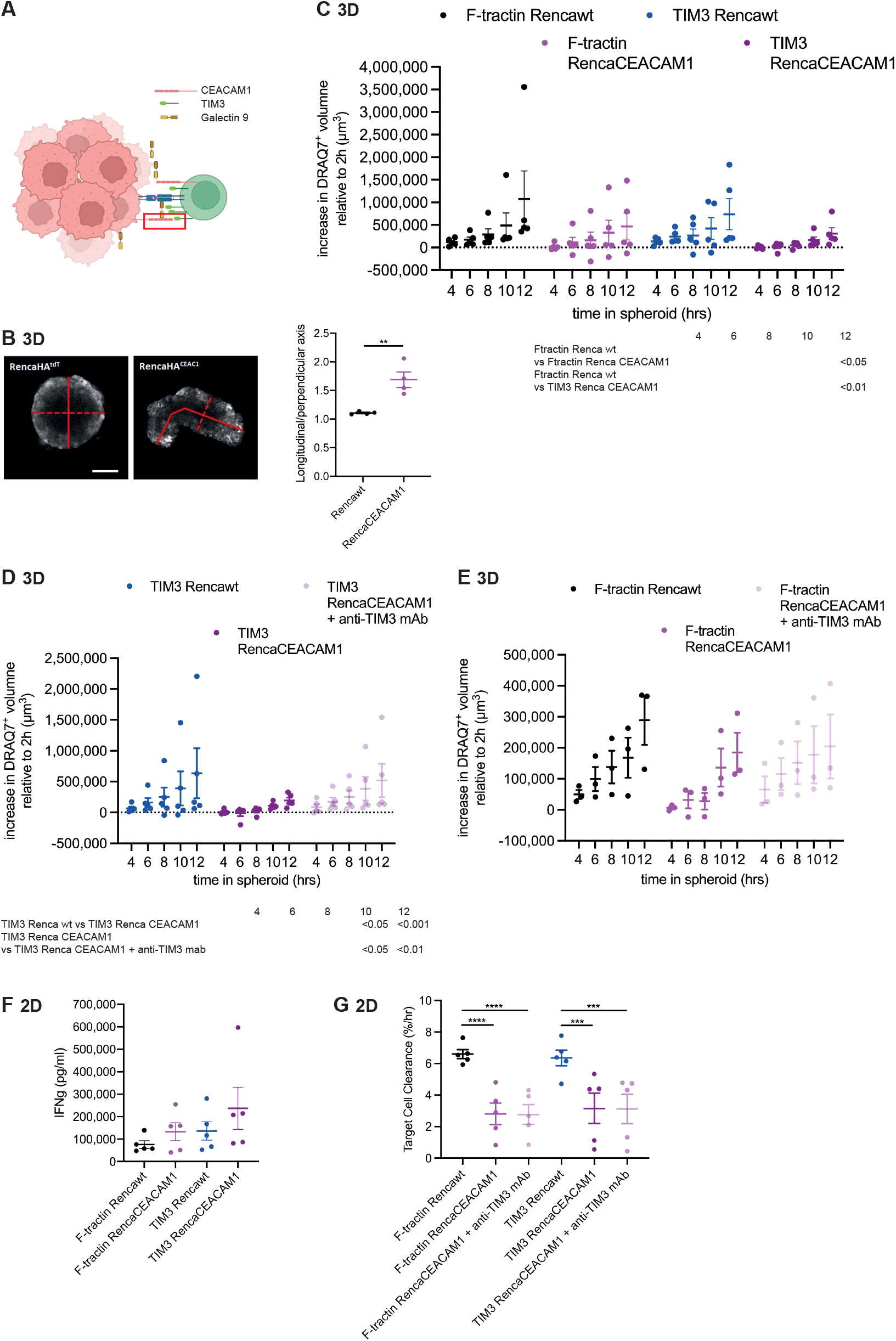
CEACAM1 in trans is an inhibitory TIM3 ligand. **A** Scheme as in Fig. 5A. Red box shows an interaction between TIM3 and CEACAM1 on Renca cells in trans. Figure created with BioRender.com. **B** Representative images of RencaHA and RencaHA CEACAM1-tdTomato spheroids with longitudinal (solid line) and perpendicular (broken line) axes indicated and the ratios of longitudinal to perpendicular axis (N=4). **C** CL4 CTL retrovirally transduced to express F-tractin-GFP or TIM3-GFP cocultured with RencaHA tdTomato or RencaHA CEACAM1-tdTomato spheroids incubated with K^d^HA peptide for 12h with images acquired every 2h. Each data point is an independent experiment (N=5) with a total of 13-15 spheroids analyzed per condition. Spheroid death, as measured by the increase in DRAQ7^+^ spheroid volume, is shown. Significance of differences between conditions at indicated time points is given in the table below. Single spheroid and spheroid infiltration data are given in Fig. S6C, D. **D, E** CL4 CTL retrovirally transduced to express TIM3-GFP (D) or F-tractin-GFP (E) cocultured with RencaHA tdTomato, RencaHA CEACAM1-tdTomato spheroids ± 10µg/ml anti-TIM3 mAb (clone RMT3-23) incubated with K^d^HA peptide for 12h with images acquired every 2h. Each data point is an independent experiment (N=5, 3) with a total of 9-14 spheroids analyzed per condition. Spheroid death, as measured by the increase in DRAQ7^+^ spheroid volume, is shown. Significance of differences between conditions at indicated time points is given in the table below. No significant differences between conditions in E. **F** IFNγ amounts in the supernatants of killing assays the end of the assay of K^d^HA-pulsed Renca tdTomato or Renca CEACAM1-tdTomato tumor cells by CL4 CTL expressing F-tractin-GFP or TIM3-GFP from 5 independent experiments. **G** *In vitro* killing assays corresponding to F with addition of conditions +10µg/ml anti-TIM3 mAb (clone RMT3-23). Average cell death rates calculated as percentage decrease in area covered by tumor cells per hour are means ± SEM. ** p < 0.01, *** p<0.001, **** p<0.0001, p values calculated using paired Students’ t-test (B) and one-way (G) and two-way (C, D) ANOVA

### Galectin9 enhances TIM3 function

Galectin9 is expressed by tumor cells and could serve as a ligand for TIM3 ^35^ (Fig. 7A). To investigate how Galectin9 expression by tumor cells regulates TIM3 function, we transfected Renca cells to express Galectin9-mCherry. Such transfection increased Galectin9 expression 2.9-fold over endogenous expression in Renca cells in tissue culture (Fig. S7A, B) and 2.3-fold over endogenous expression in Renca cells isolated from subcutaneous tumors (Fig. S7C, D). Tumor cell killing by CL4 CTL expressing TIM3-GFP but not F-tractin-GFP was substantially diminished in Renca spheroids overexpressing Galectin9 (Figs. 7B, S7E). At 12h the increase in the volume of dead tumor cells was reduced from 44,000±12,000/µm^3^ to 10,000±12,000/µm^3^ (p<0.0001) upon Galectin 9 expression in Renca spheroids interacting with CL4 CTL expressing TIM3 (p<0.01)(Fig. 7C). Tumor cell killing was restored in the presence of the TIM3 blocking mAb RMT3-23 (Fig. 7C). Spheroid infiltration was unaltered (Fig. S7F). Confirming galectin9-dependence of the inhibition of cytolysis, tumor spheroid killing was enhanced by the galectin9-blocking mAb RG9-35.7 (Fig. S7G). These data establish that Galectin9 at moderate overexpression levels serves as an inhibitory ligand for TIM3 in the three-dimensional spheroid context. When CL4 CTL expressing TIM3-GFP interacted with Renca target cells in 2D, Galectin9 overexpression by the Renca cells yielded a significant (p<0.001) increase in IFNγ generation from 130±20ng/ml to 340±45ng/ml (Fig. 7D), consistent with Galectin9 serving as a costimulatory ligand for TIM3. However, target cell killing was significantly (p<0.01) impaired upon Galectin9 overexpression (Fig. 7E). Changes in tumor cell biology are conceivable, as not further explored.

**Figure 7.**
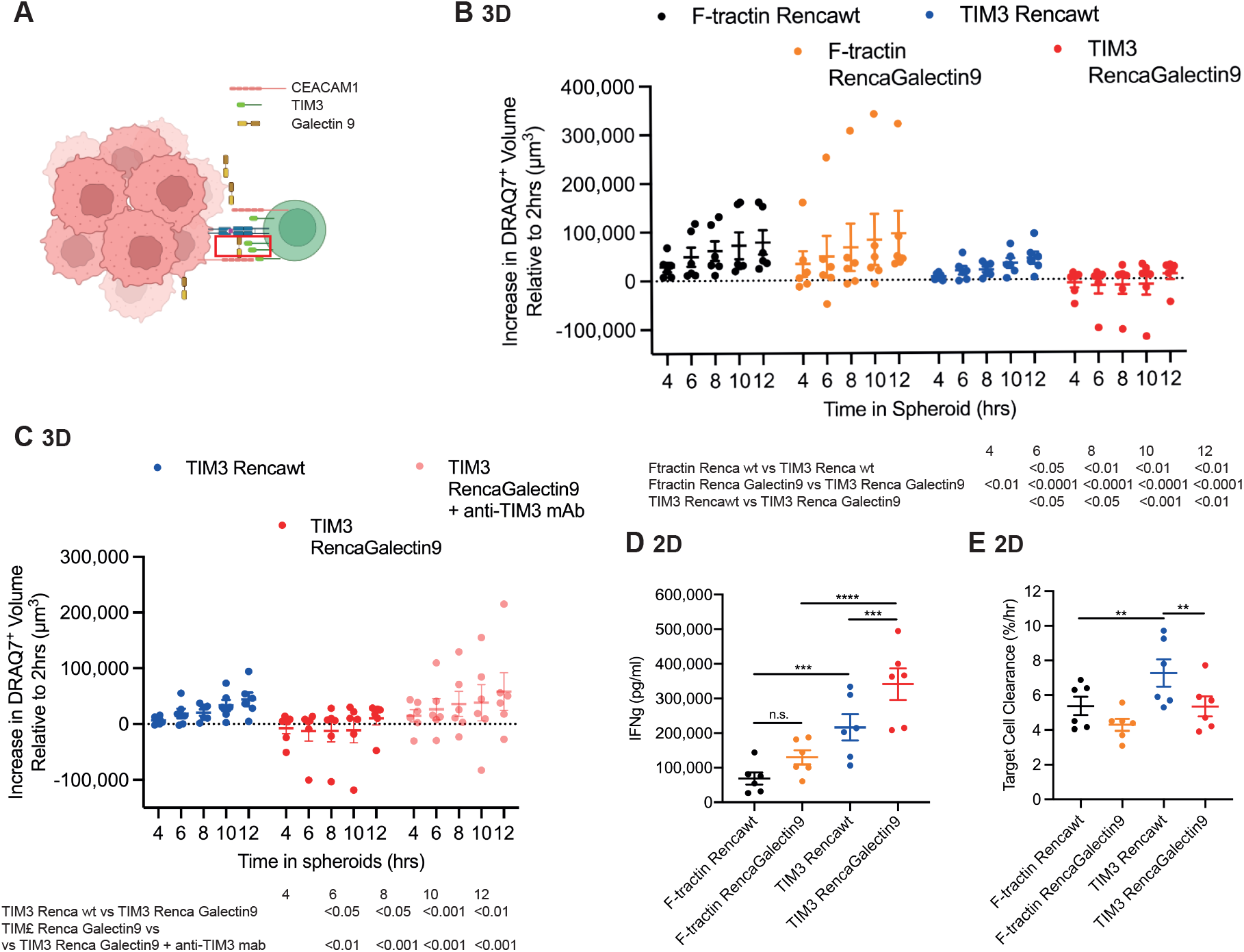
Galectin9 is an inhibitory TIM3 ligand. **A** Scheme as in Fig. 5A. Red box shows an interaction between TIM3 and Galectin9 overexpressed by Renca cells in trans. Figure created with BioRender.com. **B** CL4 CTL retrovirally transduced to express F-tractin-GFP or TIM3-GFP cocultured with RencaHA tdTomato or RencaHA Galectin9-tdTomato spheroids incubated with K^d^HA peptide for 12h with images acquired every 2h. Each data point is an independent experiment (N=6) with a total of 17-24 spheroids analyzed per condition. Spheroid death, as measured by the increase in DRAQ7^+^ spheroid volume, is shown. Significance of differences between conditions at indicated time points is given in the table below. Single spheroid and spheroid infiltration data are given in Fig. S7E, F. **C** CL4 CTL retrovirally transduced to express TIM3-GFP cocultured with RencaHA tdTomato, RencaHA Galectin-tdTomato spheroids ± 10µg/ml anti-TIM3 mAb (data without antibody the same as in B) incubated with K^d^HA peptide for 12h with images acquired every 2h. Each data point is an independent experiment (N=6) with a total of 18-21 spheroids analyzed per condition. Spheroid death, as measured by the increase in DRAQ7^+^ spheroid volume, is shown. Significance of differences between conditions at indicated time points is given in the table below. **D** IFNγ amounts in the supernatants of killing assays the end of the assay of KdHA-pulsed Renca tdTomato or Renca Galectin9-tdTomato tumor cells by CL4 CTL expressing F-tractin-GFP or TIM3-GFP from 6 independent experiments. **E** *In vitro* killing assays corresponding to D. Average cell death rates calculated as percentage decrease in area covered by tumor cells per hour are means ± SEM. ** p < 0.01, *** p<0.001, **** p<0.0001, p values calculated using one-way (D, E) and two-way (B, C) ANOVA

Together the TIM3 ligand data establish that CEACAM1 expression in cis on CTL interferes with TIM3 function, CEACAM1 and galectin9 expression in trans enhanced inhibitory TIM3 function in SIL and costimulatory TIM3 function in IFNγ secretion by CTL. Effects on 2D cytolysis were more complex.

### CTL subcellular polarization is related to cytolysis, calcium signaling to IFNγ secretion

The divergent effect of the overexpression of CEACAM1 and Galectin9 on CTL function in 2D, enhancement of IFNγ secretion but inhibition of target cell killing (Figs. 6F, 6G, 7D, 7E), offers an opportunity to determine which elements of proximal CTL activation are related to these two elements of CTL effector function. Overexpression of CEACAM1 or Galectin9 by Renca cells impaired the cytoskeletal polarization of CL4 CTL interacting with them, in particular in CTL overexpressing TIM3. While cell coupling was not substantially altered (Fig. 8A), the fraction of CL4 CTL expressing TIM3 with off-interface lamellae increased from 51±7% in contact with Rencawt cells to 81±5% and 85±5% (p<0.01) in contact with Renca cells overexpressing CEACMA1 and Galectin9, respectively (Fig. 8B). The time to the first off-interface lamella was reduced (Fig. S8A). The percentage of CL CTL4 displaying translocations remained too small to reveal substantial differences (Fig. 8C). When imaging the interaction of CL4 CTL with Renca cells overexpressing CEACAM1 or Galectin9 we noted that CL4 CTL didn’t round up as observed on Rencawt cells (Fig. 8D-H). This effect was more pronounced in CL4 CTL expressing TIM3. For example, three minutes after the formation of tight cell couples the ratio of T cell length, the axis perpendicular to the CTL tumor target cell interface, to T cell width, parallel to the interface, was significantly increased from 1.35±0.05 to 1.8±0.1 (p<0.001)(Fig. 8F) in CTL expressing TIM3 upon overexpression of CEACAM1 on the Renca cells. This inability to round up is consistent with the substantial reduction in cytolysis under the same conditions (Figs. 6G, 7E). Thus, an impaired CL4 CTL ability to polarize towards the tumor target cells was consistently related to diminished cytolysis, confirming the critical importance of such polarization for cytolysis.

**Figure 8.**
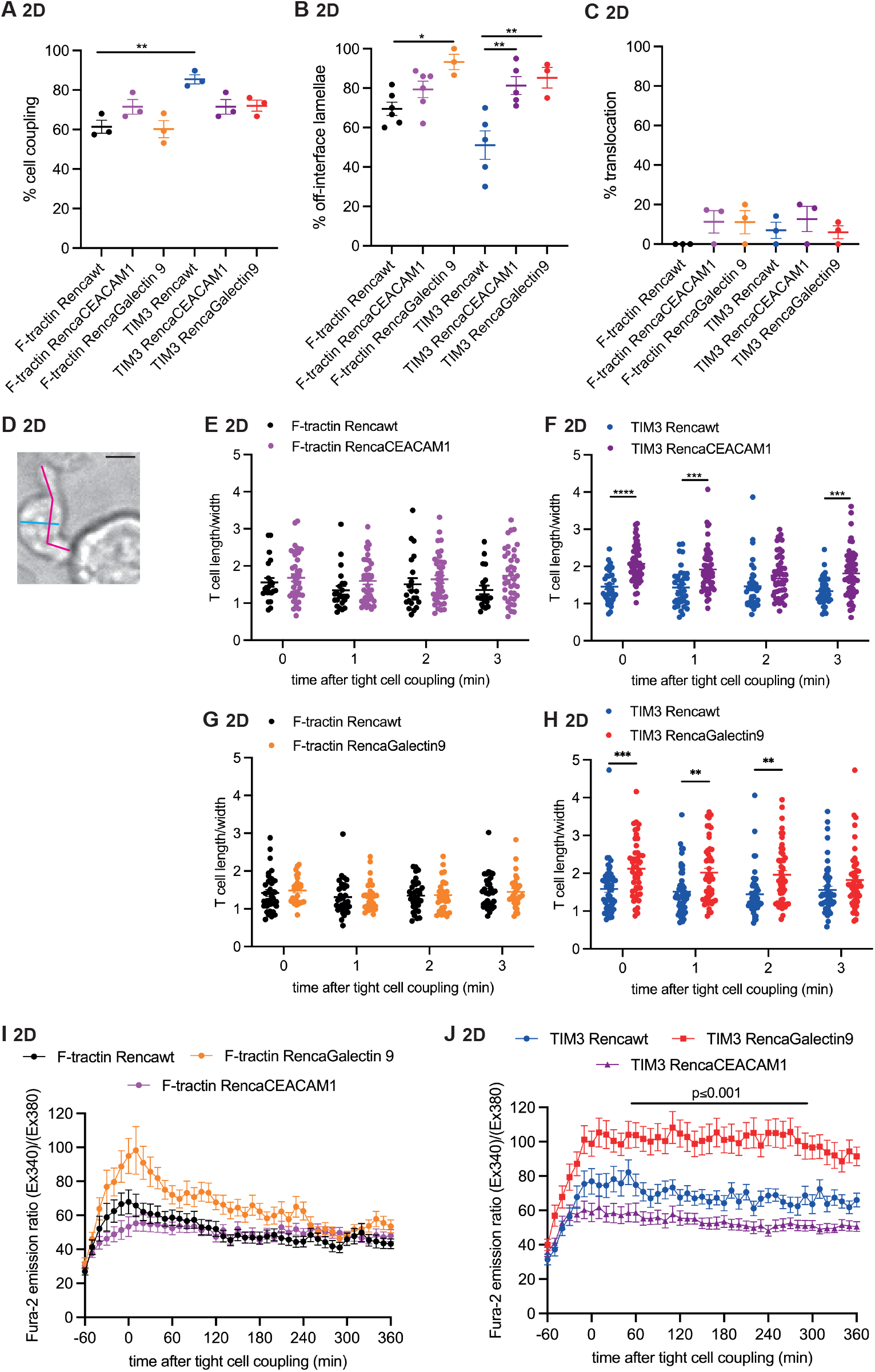
CTL morphology relates to cytolysis and calcium signaling to IFNγ secretion. **A-H** Microscopy analysis of interactions over time of Clone 4 CTL expressing F-tractin-GFP or TIM3-GFP with HA peptide pulsed Renca cells, wildtype or overexpressing CEACAM-1-tdTomato or Galectin9-mCherry. Percentage of Clone 4 CTL that form a tight cell couple upon contact with a tumor target cell (A), with off-interface lamellae (B), translocation of more than one immune synapse diameter (C). Data are means ± SEM from ≥ 3 independent experiments. Cell couples were imaged for 10±0.5min (at least 2min). D Representative imaging data showing an elongated CL4 CTL in a cell couple with a HA peptide pulsed Renca cell overexpressing CEACAM1-tdTomato 10min after tight cell coupling. Blue line is T cell width, red line is T cell length. Scale bar=5µm E, F Ratio of T cell length to width of CTL expressing F-tractin-GFP or TIM3-GFP (22-53) interacting with Rencawt or Renca-CEACAM1-tdTomato cells pooled from N=2-4 independent experiments. G, H Ratio of T cell length to width of CTL expressing F-tractin-GFP or TIM3-GFP (29-50) interacting with Rencawt or Renca-Galectin9-mCherry pooled from N=3-4 independent experiments. **I, J** CL4 CTL expressing F-tractin-GFP or TIM3-GFP interacted with Renca cells, wildtype or overexpressing CEACAM-1-tdTomato or Galectin9-mCherry, incubated with 2µg/ml K^d^HA peptide. The ratio of Fura-2 emissions at 510nm upon excitation at 340nm over 380nm is given relative to time of tight cell coupling as the mean ± SEM. N=2-3 independent experiments, 39-50 T cells analyzed per condition. Single cell and individual experiment average data are given in Fig. S8B, C. * p<0.05, ** p < 0.01, *** p<0.001, **** p<0.0001, p values calculated using one-way (A-C, J for each time point) and two-way (E – H, data log transformed) ANOVA

In contrast to diminished CTL polarization, overexpression of CEACAM1 or Galectin9 on Renca cells yielded a consistent increase in calcium signaling in CL4 CTL interacting with these tumor target cells (Figs. 8I, J, S8B, C). This effect was more pronounced upon overexpression of Galectin9 on the Renca cells and TIM3 in the CL4 CTL. For example, the emission ratio of the calcium-sensitive dye Fura-2, a measure of calcium signaling, was almost doubled upon Galectin9 expression in CTL expressing TIM3 (p<0.001 across the majority of time points)(Fig. 8J). Such elevated calcium signaling closely mirrored IFNγ secretion by the CL4 CTL (Figs. 6F, 7D), suggesting critical importance of calcium signaling for IFNγ secretion. Our data thus relate different proximal signaling processes to divergent CTL effector function, cytoskeletal polarization to cytolysis and calcium signaling to cytokine secretion.

### The physiological environment for TIM3 function is more inhibitory in spheroids

To further understand the inhibitory role of TIM3 selectively in the 3D spheroid environment, we determined molecular features of this environment and potential consequences for CL4 SIL and CL4 CTL function. As an initial control to determine whether CL4 CTL contact with the Matrigel extracellular matrix, part of the tumor cell spheroids, in itself could induce TIM3 inhibitory function in CL4 CTL, we cultured CL4 CTL in Matrigel overnight. In contrast to co-culture with tumor cells in spheroids (Fig. 1E), TIM3 expression did not alter the ability of CL4 CTL to kill RencaHA tumor target cells after Matrigel only culture (Fig. 9A). These data establish that 3D tumor cell contact was required for the inhibitory function of TIM3.

**Fig. 9.**
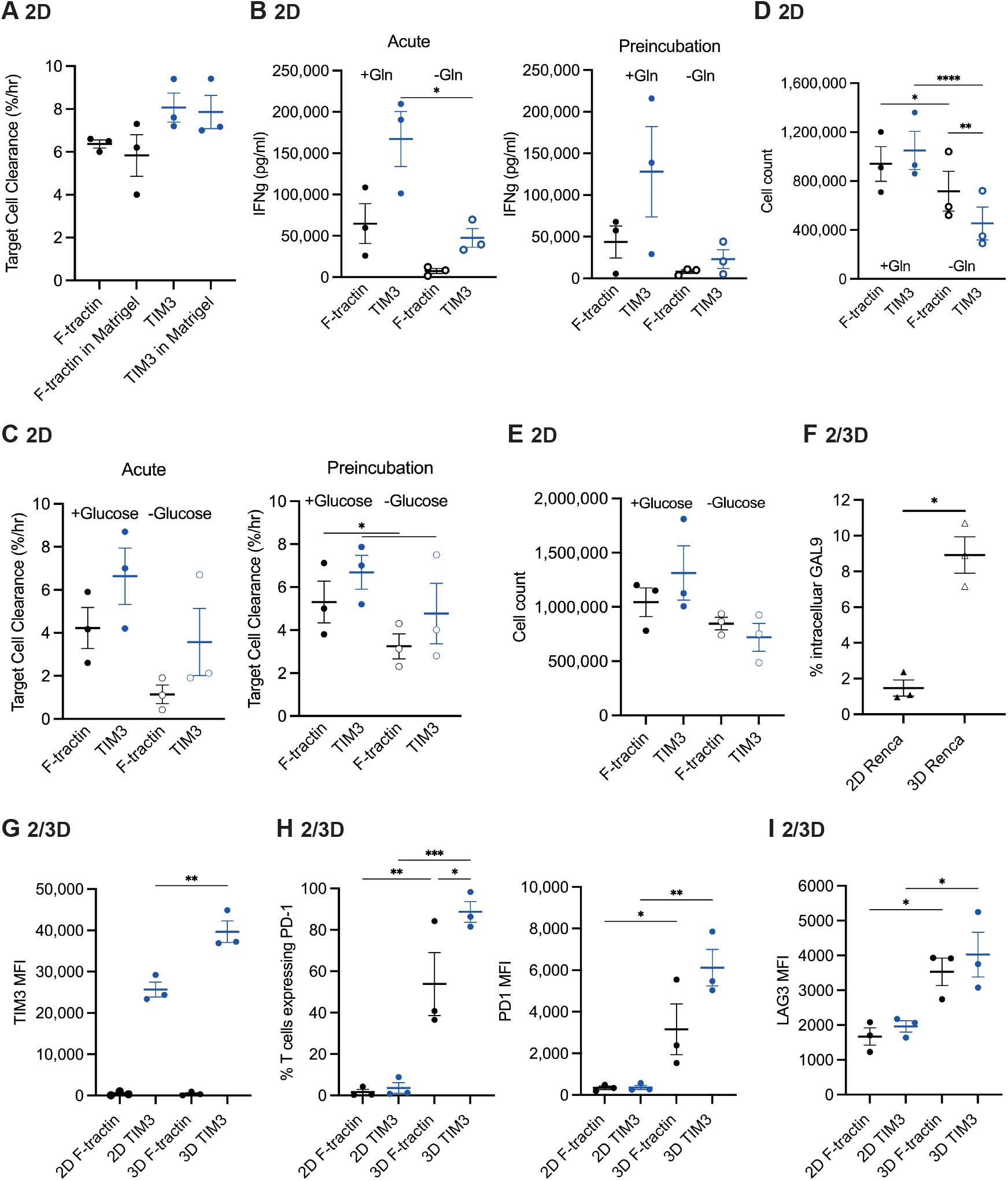
The physiological environment for TIM3 function is more inhibitory in spheroids. **A** *In vitro* killing of KdHA-pulsed Renca tumor cells by CL4 CTL expressing F-tractin-GFP or TIM3-GFP after overnight culture in Matrigel or medium only. Average cell death rates calculated as percentage decrease in area covered by tumor cells per hour are means ± SEM from 3 independent experiments. **B-E** CL4 CTL were assayed in the absence of glutamine (‘Gln’) or glucose (‘Acute’) or incubated overnight without glutamine or glucose and assayed thereafter in the presence of the nutrients (‘Preincubation’). B IFNγ amounts in the supernatant of killing assays of *in vitro* killing of KdHA-pulsed Renca tumor cells by CL4 CTL expressing F-tractin-GFP or TIM3-GFP ± glutamine. 3 independent experiments. Killing assays in Fig. S9A. C *In vitro* killing of KdHA-pulsed Renca tumor cells by CL4 CTL expressing F-tractin-GFP or TIM3-GFP ± glucose. 3 independent experiments. IFNγ amounts in the supernatant in Fig. S9B. D, E Number of CL4 CTL after overnight culture ± glutamine (D) or glucose (E). 3 independent experiments each. **F** Percentage RencaHA cells with high intracellular galectin9 after overnight culture in 2D or spheroids (‘3D’). 3 independent experiments. Representative staining data in Fig. S9C. **G-I** Mean fluorescence intensity (MFI) of staining and percent T cells positive for TIM3 (G), PD-1 (H) and LAG3 (I) as indicated of CL4 CTL expressing F-tractin-GFP or TIM3-GFP after overnight culture with 3D spheroids (‘3D’) or 2D culture with IL-2 (‘2D’). % T cells positive for TIM3 and LAG3 are in Fig. S9F. 3 independent experiments. * p<0.05, ** p < 0.01, **** p<0.0001, p values calculated using one-way ANOVA (B -D, G – I) and paired student’s t-test (F).

Tumor cells compete with CTL for nutrients in the tumor microenvironment ^36^. To determine whether nutrient limitation could switch TIM3 from the costimulatory function in 2D tissue culture (Fig. 1F, G) to inhibitory function (Fig. 1C), we determined CL4 target cell killing and IFNγ secretion in two 2D settings of nutrient depletion. We either withheld glutamine or glucose during the killing assay (‘acute’) or we cultured CL4 CTL in the absence of glutamine or glucose overnight to mimic adaptation to an extended presence in a nutrient-limiting environment (‘preincubation’). While glutamine or glucose were required for effective CL4 IFNγ secretion and killing, respectively, TIM3 still functioned as a costimulatory receptor under these nutrient-deprived conditions (Figs. 9B, C, S9A, B). However, CL4 CTL expansion during the overnight incubation in the absence of glutamine or glucose was reduced (Fig. 9D, E). While under the nutrient-replete control conditions TIM3 functioned as a costimulator of CL4 CTL proliferation as expected, in the absence of glutamine, and less prominently glucose, TIM3 switched to inhibitory function, reducing CL4 CTL expansion overnight (Fig. 9D, E). These data suggest that metabolic restriction supports some but not all elements of TIM3 function as an inhibitory receptor.

Next, we determined the expression of TIM3 ligands on the RencaHA tumor target cells grown in spheroids versus 2D. After spheroid culture, the size of a population of RencaHA cells with Galectin9 expression elevated by more than a log was significantly (p<0.05) increased from 1.5±0.5% to 9±1% (Figs. 9F, S9C). Expression of CEACAM1 was minimal under both conditions (Fig. S9D, E). Parallel expression of inhibitory receptors on CL4 CTL could alter the signaling context of TIM3 function. Expression of TIM3-GFP itself in CL4 SIL was significantly (p<0.01) elevated from a mean fluorescence intensity of 25,500±3,000 in CL4 CTL to 39,500±4,500 (Fig. 9G). This was despite sorting CL4 T cells for the same levels of TIM3-GFP expression before the start of the assay. Expression of PD-1 and LAG3 was also elevated in CL4 SIL (Figs. 9H, I, S9F) but not that of CD39 (Fig. S9G).

In combination these data establish that the molecular environment for TIM3 function is more inhibitory in tumor cell spheroids: expression of multiple inhibitory receptors was elevated on the CL4 SIL; potential limitation of glutamine and glucose could turn TIM3 inhibitory in T cell proliferation. In addition, enhanced expression of TIM3 itself and of galectin9 as a TIM3 ligand in spheroids may lead to stronger TIM3 signaling. In combination all data presented here suggest that TIM3 functions as a signal amplifying co-regulatory receptor. In the inhibitory environment of spheroids TIM3 would function as a coinhibitor, in the stimulatory environment of 2D tissue culture as a costimulator.

## Discussion

### TIM3 directly regulates cytolysis of tumor target cells

Here we have used primary murine and human CTL and tumor cell spheroids ^12^ to investigate TIM3 function in the direct interaction of CTL with tumor target cells. We have virally expressed TIM3 in CL4 CTL as TIM3-GFP at levels similar to those of endogenous TIM3 in CTL stimulated repeatedly with persistent antigen. We have expressed TIM3 in human CTL at a slightly lower level (Fig. S2A-C). At those levels, TIM3 directly suppressed the killing of tumor cells by CTL in spheroids (Figs. 1, 2). The regulation of tumor cell killing by CTL complements roles of TIM3 in regulating chemokine secretion and the suppression of innate immunity to double stranded DNA by dendritic cells, in the uptake of apoptotic debris and the sensitization of CTL to fratricide by macrophages and in cytokine secretion by and apoptosis of Th1 CD4^+^ T cells ^10, 13, 14, 15, 16, 37, 38^. Blocking TIM3 with an antibody could fully restore tumor target cell killing in our spheroid system (Fig. 1C). Yet, given the multiple other roles of TIM3 it remains uncertain whether the enhancement of tumor cell killing by TIM3 blockade would be the dominant effect of anti-TIM3 treatment in vivo. Nevertheless, the direct regulation of CTL killing by TIM3 contrasts with roles of PD-1 and CTLA-4. Using the same CL4 CTL, direct tumor cell killing by CL4 CTL is not re-activated by blocking PD-1 ^12^ and myeloid cell-supported clonal T cell replacement offers an alternate explanation for the efficacy of in vivo anti-PD-1 treatment ^4, 39^. The CTLA-4 ligands CD80 and CD86 are not commonly expressed on tumor cells. Given these differences in mechanisms of action, blocking TIM3 promises to be synergistic with PD-1 and/or CTLA-4 blockade as already explored in early-stage clinical trials ^7, 40^.

### CEACAM1 and galectin9 are TIM3 ligands in the regulation of CTL function

Our reductionist approach to CTL suppression allowed us to identify TIM3 ligands in the regulation of tumor cell cytolysis. Galectin9 expressed by tumor cells enhanced the inhibitory function of TIM3 (Fig. 7B). This enhancement could be reversed by the antibody RMT3-23 (Fig. 7C). As this antibody does not interfere with the TIM3-Ig galectin9 interaction in an ELISA ^18^, the effect of galectin9 is likely indirect. As galectin9 is a glycan-binding protein, it can interact with multiple receptors at the interface between CTL and tumor target cells ^21^, including an already described interaction with PD-1 ^20, 41^. CEACAM1 on tumor cells functioned as a second inhibitory ligand for TIM3 (Fig. 6B). Consistent with a contribution of a direct interaction, this inhibitory effect was reversed by blockade with the anti-TIM3 antibody RMT3-23 (Fig. 6C) that blocks the CEACAM1 TIM3 interaction ^18^. Nevertheless, indirect effects are also likely as CEACAM1 expression altered tumor spheroid morphology (Fig. 6B) and as effects of CEACAM1 expression by tumor cells on CTL-mediated cytolysis in a conventional 2D tissue culture system could not be blocked by RMT3-23 (Fig. 6G). CEACAM1 expression on CL4 CTL in cis interfered with the inhibitory function of TIM3 (Fig. 4B). One possible explanation is altered TIM3 trafficking. CEACAM1 co-expression reduced TIM3 cell surface expression by almost 10-fold even though overall cellular TIM3 levels were reduced much less (Fig. S6A, B), consistent with diminished trafficking to the plasma membrane and/or enhanced TIM3 endocytosis. An alternate explanation for the inhibitory effect of CEACAM1 co-expression is blockade of TIM3 ligand engagement by competition with direct binding between the CEACAM1 and TIM3 N-terminal Ig domains ^17^. A previous study found that CEACAM1 was required for effective TIM3 expression ^17^. As our use of retroviral expression of TIM3 possibly bypasses a need for CEACAM1 in elevating TIM3 expression, we could isolate an inhibitory effect of cis CEACAM1 expression on TIM3 function here.

### TIM3 is a context-dependent co-regulatory receptor

While TIM3 expression inhibited cytolysis in tumor cell spheroids, using the same Renca tumor cells and CL4 CTL, TIM3 expression enhanced cytolysis and IFNγ secretion in conventional 2D tissue culture (Fig. 1). This costimulatory effect is consistent with enhanced tyrosine phosphorylation, Akt and MAPK activation and increased NFAT and NFκB nuclear translocation observed upon TIM3 expression ^5, 23, 24, 42^. However, the TIM3 signalosome also contains inhibitory signaling intermediates such as SHP-1, Cbl-b and UBASH3A ^25^. Moreover, Lck as a key TIM3-associated signaling intermediate ^21, 25, 38, 43^ mediates signaling by antigen receptors as well as inhibitory receptors ^44^. Given this dual costimulatory and coinhibitory potential of TIM3, signaling effects of TIM3 engagement may depend on other signaling pathways currently active in the cell. In CL4 CTL interacting with spheroids, expression of the inhibitory receptors PD-1 and LAG3 was elevated (Fig. 9I, J). Moreover, upon glucose and glutamine limitation, as likely experienced in CTL competition with tumor cells ^36^, TIM3 lost its ability to costimulate CTL proliferation (Fig. 9C, E). We, therefore, suggest that TIM3 functions as a context-dependent signal amplifying co-regulator. When inhibitory signaling pathways are engaged in a T cell, TIM3 will enhance such signaling to function as a coinhibitory receptor. When activating signaling dominates, e.g. during strong TCR engagement in the presence of abundant nutrients as commonly used in signaling studies, TIM3 will amplify such signaling to function as a costimulatory receptor. As in vivo TIM3 is highly expressed on CTL in the inhibitory setting of persistent antigen exposure, its coinhibitory role can be expected to dominate.

### Context-dependent elements of TIM3 signaling

Tim3 signals through association with other receptors, translocation to the cellular interface with an interacting cell as mediated by the transmembrane domain, and through signaling motifs in the cytoplasmic domain, prominently phosphorylation of tyrosine residues 256 and 263 ^23, 24, 25^. TIM3 can regulate the nuclear translocation of the transcription factors NFAT and NFkB, T cell apoptosis via Bat3 and glucose metabolism ^23, 24, 38, 45^. Here we have shown that a key mechanism of TIM3 action in CTL, shared between coinhibitory and costimulatory function, is the regulation of cytoskeletal polarization. A key element of CTL suppression is the inability to maintain a CTL tumor cell couple ^11, 12^. TIM3 expression further impaired the ability of CL4 SIL to maintain cell couples (Fig. 3E). Analogously, costimulatory TIM3 signaling in 2D tissue culture further stabilized CL4 CTL tumor cell couples (Fig. 3B, C). As a potential mediator of this effect, the established cytoskeletal regulator Vav1 ^46, 47^ is part of the TIM3 interactome ^25^. However, clear distinctions between coinhibitory and costimulatory TIM3 signaling also emerged. Inhibitory signaling of TIM3 in spheroids could the fully blocked with the antibody RMT3-23 (Fig. 1C), suggesting that it is fully dependent on ligand engagement. TIM3 lacking the cytoplasmic domain was substantially less inhibitory in this context (Fig. 4A), suggesting a direct role for signal transduction through motifs in the TIM3 cytoplasmic domain. In contrast, costimulatory signaling of TIM3 in 2D tissue culture was only partially blocked with RMT3-23 (Fig. 1F, G) with an only modest effect of the lack of the TIM3 cytoplasmic domain (Fig. 4B, C). In addition, the inhibitory role of TIM3 in spheroids was apparent both at the high and more modest expression levels of TIM3 in murine and human CTL, respectively (Figs. 1C, 2B). In contrast, the costimulatory role of TIM3 was limited to the higher expressing murine CTL (Figs. 1F, G, 2C, D). These data suggest a mechanistic difference between TIM3 inhibitory and stimulatory signaling, the former largely dependent on ligand engagement and cytoplasmic association of signaling intermediates, the latter more likely to also involve alternate mechanisms, possibly through lateral association with other receptors such as CD2 and OX40 as parts of the TIM3 interactome ^25^.

Our investigation of TIM3 in the 2D costimulatory context allowed us to determine the association of signaling processes with cytolysis versus IFNγ secretion. While both are enhanced upon simple expression of TIM3 in CL4 CTL (Fig. 1F, G), combining such expression with expression of CEACAM1 and galectin9 on the tumor cells led to divergent effects, cytolysis was suppressed while IFNγ secretion was enhanced (Figs. 6F, 6G, 7D, 7E). Divergent regulation of cytolysis and IFNγ secretion has been described before ^48, 49^, with diminished F-actin generation upon HEM1 deficiency linked to enhanced lytic granule release and reduced cytokine secretion^50^. Here we have shown that cytolysis was consistently associated with the ability of the CTL to maintain a stable cell couple (Fig. 8A, B), i.e. cytoskeletal regulation. In contrast, IFNγ secretion was consistently associated with calcium signaling (Fig. 8I, J), possibly through control of NFAT-dependent transcriptional regulation. While cytoskeletal polarization and calcium signaling have been linked ^51^, our data suggest that such linkage is only partial.

## Acknowledgments

We acknowledge support from the University of Bristol Flow Cytometry and Wolfson BioImaging core facilities. This work was supported by grants from the MRC (MR/N0137941/1 to CCWW and MR/W006308/1 to TG for the GW4 BIOMED MRC DTP), the Wellcome Trust (201254/Z/16/Z/ to GLE), Cancer Research UK (DRCRPG-NOV21/100003 to AMG) and Immunocore (to CJH and CW). HA, AA and SA were supported by the Ministry of Education of Saudi Arabia.

## Author contributions

HA and CCWW designed and carried out experiments, analyzed data and wrote the paper. AA, GLE, TG, SA, JB carried out experiments and analyzed data. CJH provided reagents and reviewed and edited the paper. DJM and AMG provided supervision and reviewed and edited the paper. CW conceptualized the study, designed and carried out experiments, analyzed data and wrote and edited the paper.

## Methods

### Mice

Thy1.2^+/+^ BALB/c, (Charles River, Oxford, UK, Research Resource Identifier (RRID): IMSR_CRL547) and Thy1.1^+/+^ CL4 TCR-transgenic mice (RRID: IMSR_JAX:0053079) were maintained at the University of Bristol Animal Services Unit. All mouse experiments were compliant with UK Home Office Guidelines under PPL 30/3024 to DJM as reviewed by the University of Bristol AWERB (Animal welfare and ethical review body) committee.

### Human blood samples

Blood buffy coats from anonymous donors were purchased from NHS-BT with human work approved by the London-Riverside Research Ethics Committee under reference number 20/PR/0763.

### Antibodies

Antibodies used are described in the order: antigen, fluorescent label, clone, supplier, dilution/concentration, RRID:

For flow cytometry:

Murine TIM3, PE, B8.2C1.2, Biolegend, 1:200, RRID: AB_1626181

Human TIM3, PE, F38-2E2, eBioscience,1:20, RRID: AB_2572605

Human TIM3, SB600, F38-2E2, eBioscience,1:20, RRID: AB_2688208

Murine galectin9, APC, REA1069, Miltenyi Biotec, 1:30, RRID: AB_2733298

Murine CEACAM1, APC, CC1, Biolegend, 1:100, RRID: AB_2632612

Murine PD-1, BV785, 29F.1A12, Biolegend, 1:200, RRID: AB_2563680

Murine LAG3, Pe-Cy7, C9B7W, eBioscience, 1:200, RRID: AB_2573428

Murine CD39, PerCP-eFluor710, 24DMSI, eBioscience, 1:100, RRID: AB_10717953

For blocking:

Murine TIM3 no azide (for *in vitro/in vivo* blockade) RMT3-23 BioXcell In Vivo mAb *in vivo:* 100μg/mouse in vitro: 10μg/ml RRID: AB_10949464

Isotype control for anti-TIM3 Rat IgG2a 2A3 no azide (for *in vivo/in vitro* blockade) BioXcell In Vivo mAb *in vivo:* 100μg/mouse *in vitro:* 10μg/ml RRID: AB_1107769

Murine galectin9 pre-functional grade, RG9-35.7, Miltenyi Biotec, 10 µg/ml. RRID: AB_2651797

### Murine cell culture

The murine Renal Carcinoma cell line Renca (RRID: CVCL_2174) and the retrovirus-producing cell line Phoenix (RRID: CVCL_H717) were maintained as previously described ^12^.

To generate *in vitro* CL4 CTL, CL4 mouse spleens from 6-12-week-old male or female mice were macerated. Red blood cells were lysed using ACK Lysis buffer (Gibco, Thermo Fisher), and the remaining splenocytes were resuspended in RPMI-1640 complete medium with 10% FBS, 2mM Glutamine and 50µM 2-mercaptoethanol (RPMI complete medium). 5×10^6^ cells were seeded into each flat bottomed 24 well tissue culture plate with 1μg/ml of K^d^HA peptide (IYSTVASSL^[518-526]^) from influenza virus A/PR/8/H1N1, for 24h at 37°C. After 24h, cells were washed 5 times in RPMI (Gibco) and reseeded in to 24 well pates at 5×10^6^ cells per well in 2ml RPMI complete medium containing 50 units/ml of recombinant human IL-2 (National Institutes of Health/NCI BRB Preclinical Repository). Retroviral transduction was performed if required as previously described ^12^. CL4 T cells were then passaged using the same IL-2 containing medium.

### Human cell culture

Human HepG2 hepatoblastoma (RRID: CVCL_0027) and Mel624 melanoma (RRID: CVCL_8054) cells were stably transfected to express mCherry or tdTomato ^52^. HepG2 cells were maintained in RPMI complete medium. Mel624 cells were maintained in high glucose DMEM with 10% FBS, 2mM Glutamine, 1mM pyruvate (DMEM complete medium). Medium for the growth of mCherry or tdTomato transfectants was supplemented with 250μg/ml Hygromycin. Blood buffy coats were obtained from healthy donors. Peripheral Blood Mononuclear cells (PBMC) were isolated by density gradient centrifugation using Ficoll-Paque^TM^ (Sigma-Aldrich). PMBCs were cryopreserved at a concentration of 2.5×10^7^ cells/ml. To isolate CD8^+^ T cells, buffy coat cryopreserved vials were thawed, and the cells were washed twice with RPMI complete medium. Cells were resuspended in ice cold MACS buffer. CD8^+^ T cells were purified by magnetic enrichment for CD8^+^ cells using CD8 MicroBeads (Miltenyi) following the manufacturer’s protocol. CD8^+^ T cells were activated using CD3/CD28 Dynabeads (Life Technologies) at bead-to-cell ratio of 1:1 in human IL-2 medium (X-VIVO 15, serum-free hematopoietic cell medium, with 2mM L-Glutamine and gentamicin (Lonza) supplemented with 5% Human AB serum (Valley Medical), 10mM neutralized N-acetyl L-Cysteine (Sigma-Aldrich), 50μM β-Mercaptoethanol (Gibco, Thermo Fisher), and 30U/ml rh-IL-2 (NIH/NCI BRB Preclinical Repository) – human IL-2 medium) and incubated overnight at 37 °C and 6% CO2. The 1G4 TCR ^53^ was expressed in primary human T cells using a pHR_SFFV -based lentiviral vector (RRID_Addgene79121) with an expression cassette of alpha chain-P2A-beta chain-P2A-GFP/F-tractin-GFP/hTIM3-GFP. For lentiviral transduction, HEK 293T cells (RRID:CVCL_0063; Lenti-X 293T cells, Takara) were maintained in DMEM complete medium. For transduction 1.5×10^6^ Lenti-X 293T cells were seeded in 5ml DMEM complete medium in 60×15 mm Primaria culture plates (Corning) 24h before transfection. Cells were transfected with a total of 4.5 µg plasmid DNA using Fugene HD (Promega): 0.25µg envelope vector pMD2.G (RRID:Addgene_12259), 2µg of packaging plasmid pCMV-dR8.91 (Creative Biogene), and 2.25µg of the pHR_SFFV-based transfer vector. 48h after transfection, virus containing medium was collected and filtered through a 0.45µm nylon filter. After 24h of setting up the primary human T cell culture, 1×10^6^ T cells were mixed with 500-700µl lentivirus-containing medium in a 24-well plate medium in presence of 8µg/ml Polybrene (Sigma-Aldrich) and centrifuged for 1.5h at 2500rpm, 37 °C. After spinduction primary CD8^+^ T cells were resuspended in human IL-2 medium. Cells were maintained at density of less than 2×10^6^ cells/ml and if necessary spilt back to a density of 0.5-1×10^6^ cells/ml.

### Tumor Growth and Treatment Experiments

6-week-old Thy1.2^+/+^ BALB/c mice were injected subcutaneously, in the dorsal neck region, with 1×10^6^ RencaHA tumor cells in 100μl PBS. Tumor measurements and treatment commenced at day 12, when tumors of approximately 5 x 5mm diameter were palpable. Tumor-bearing mice were injected i.v. at day 12 with 5×10^6^ day 5 CL4 CTL (see above). Control mice received 100μg/mouse isotype control (Rat IgG2a, 2A3, BioXcell InVivoMAb)^54^. Treated mice received 100μg/mouse anti-TIM3 mAb (RMT3-23, BioXcell InVivoMAb) injected intraperitoneally in 100μl PBS on alternate days throughout the experiment. Tumors were measured on alternate days using calipers and the volume calculated using the modified elliptical formula: Volume = 0.5 x length x width^2^.

### Imaging and Image Analysis

For live cell imaging of immune synapse formation and CL4 T cell morphology 1×10^6^ Renca tumor target cells were pulsed with 2µg/ml K^d^HA for 1h at 37°C. Cells were then resuspended at 1×10^6^/400μl Imaging Buffer (PBS, 10% FBS, 1mM CaCl^2^, 0.5mM MgCl^2^).

For imaging of cytoplasmic calcium, T cells were loaded with 2µM Fura-2 (Molecular Probes) for 30min at room temperature as established ^12^. For image acquisition, 40,000 Clone 4 CTL or spheroid-infiltrating lymphocytes in 5μl imaging buffer were plated with 1-1.5μl Renca target cells (preceding paragraph) in 50μl imaging buffer, in a 384-well, glass-bottomed imaging plate (Brooks). Every 10s for 15min, one bright-field differential interference contrast (DIC) image, one fluorescence image with excitation at 340 nm, and one fluorescence image with excitation at 380nm were acquired at 37°C with a 40x oil objective (NA = 1.25) on a Leica DM IRBE-based wide-field system equipped with Sutter DG5 illumination and a Photometrics Coolsnap HQ2 camera.

Using Fiji/ImageJ ^55, 56^ for analysis of DIC images, tight cell couple formation was defined as the first time point at which a maximally spread immune synapse formed, or 40s after initial cell contact, whichever occurred first. To assess CTL and TIL morphology in cell couples with tumor target cells, every DIC frame after tight cell couple formation was assessed for the presence of off-synapse lamellae, defined as transient membrane protrusions pointing away from the immune synapse, followed by retraction. To determine CTL translocation over the Renca cell surface, the position of the immune synapse on the RencaHA target cell was compared to the position at cell coupling. If the T cell had migrated by a distance greater than the diameter of the immune synapse, this was classed as translocation. For calcium imaging analysis, rolling ball background fluorescence was subtracted from the 340nm and 380nm excitation fluorescence data, and the ratio of the Fura-2 images upon excitation at 340 versus 380 nm was determined in a circular region of interest of the dimensions of the T cell.

### Cytolysis and IFNγ secretion

For microscope-based Cytotoxicity Assays, the IncuCyte™ Live Cell analysis system and IncuCyte™ ZOOM software (Essen Bioscience) were used to quantify target cell death. 1×10^6^ Renca, MEL624 or HepG2 cells transfected to express the fluorescent protein mCherry or tdTomato were either untreated (control) or pulsed with 2μg/ml K^d^HA or HLA-A*02:01 NY-ESO-1^157-165^ (SLLMWITQC) peptide for 1h for mouse and human cells, respectively. Cells were suspended in 5ml Fluorobrite medium (Thermo Fisher) with 10% FBS, 2mM L-Glutamine, 50µM 2-mercaptoethanol to a concentration of 15,000 cells/50μl. Cells were plated in each well of a 384 well Perkin-Elmer plastic-bottomed plate and incubated for 4h to adhere. 10,000/40,000 murine/human CTL that had been FACS sorted for expression of GFP-tagged proteins were added per well to the plate in 50μl Fluorobrite medium, yielding a 1:1/4:1 effector to target ratio, respectively. Images were taken every 15min for 14h at 1600ms exposure using a 10x lens. The total red object (mCherry target cell) area (µm^2^/well) was quantified at each time point. The CTL killing rate was determined as the linear gradient of the red object data at its steepest part for 6h. The CTL killing rate was calculated as the difference to the growth rate of Renca (control) cells which were not pulsed with cognate HA antigen during the same 6h time window.

To determine IFNγ secretion, supernatants at the end of the imaging-based cytolysis assays were collected and frozen down. To measure IFNγ concentrations, mouse and human IFNγ OptEIA ELISA Kits (BD Biosciences) were used according to manufacturer’s instructions. Briefly, wells of a 96-well Maxisorb Nunc-Immuno plate (Theromfisher) were coated with 100µl anti-mouse or anti-human IFNγ monoclonal antibodies diluted in 0.1M Na^2^CO^3^ coating buffer at a 1:250 dilution and incubated overnight at 4°C. On the second day, plates were washed 5 times with ELISA wash buffer and blocked using PBS with 10% FBS for 1h. Samples were diluted in ELISA dilution buffer at 1:200 dilution for mouse CTL and 1:20 for human CTL killing assay supernatants and incubated for 1h at room temperature. Plates are washed 5 times with wash buffer. 100µl of working detectors (Biotinylated anti-mouse or anti-human IFN-ƴ monoclonal antibody + Streptavidin-horseradish peroxidase conjugate) was added to each well and incubated for 1h at RT. Then, the plate was washed 10 times with ELISA wash buffer. 100 µl of Substrate solution (BD biosciences) was added to each well and incubated for 30min at RT. 50µl/well Stop Solution (1 M H^2^SO^4^) was added. Absorbance at 450nm and 570nm were measured within 20min after addition of stop solution. Each sample was measured in duplicate or triplicate.

### Spheroids

RencaHA tdTomato or Mel624 tdTomato cells were resuspended at a concentration of 1×10^5^ cells/ml, mixed with Matrigel (Corning) at 4°C, seeded in a 24-well plate at a final concentration of 500 cells per Matrigel dome, and left to solidify for 10min at 37°C. 2ml cell medium was added to each well and cells incubated at 37°C for 11 days. Each Matrigel dome was washed twice in PBS and incubated for 1h with 1ml of Cell Recovery Solution (Corning). Spheroids were collected in a 15-ml Falcon tube and pulsed with K^d^HA or NY-ESO-1 peptide at a final concentration of 2μg/ml for 1h. Pulsed spheroids were re-embedded in Matrigel together with 5×10^6^ primed CL4 CTL per Matrigel dome. Matrigel domes were dissolved for analysis of spheroid-infiltrating T cells after 16h: Spheroids were washed twice in PBS and incubated with 1ml of Cell Recovery Solution (Corning). Spheroids were collected, washed through a 40μm sieve and then disaggregated to retrieve T cells in 500μl of imaging buffer for immediate FACS sorting.

### Spheroid imaging

Spheroids were grown as described in the preceding paragraph. On day 10, CL4 or 1G4 CTL that had been retrovirally transduced to express the GFP-tagged protein of interest (TIM3-GFP or F-tractin-GFP) were sorted by flow cytometry and incubated in IL-2 medium for 1h ± 10µg/ml anti-TIM3 mAb (Clone RMT3-23) where appropriate. Meanwhile, spheroids were dissociated from Matrigel and resuspended into fresh Matrigel at a concentration of ∼8 spheroids/µl. 50µl of the spheroid-Matrigel suspension was separated into Eppendorf tubes for each treatment group, followed by the addition of 200,000/500,000 mouse/human sorted CTL per tube. 50µl of Matrigel, containing spheroids and T cells, was plated into each well of a 24-well tissue culture plate. After Matrigel had set, 1ml of Fluorobrite medium was added to each well, containing 1.5µM DRAQ7 viability dye ± 10µg/ml anti-TIM3 mAb. Images were acquired every 2h post-plating CTL with spheroids in 3µm z steps from the bottom of the spheroid to its widest point, usually 40 steps, for 12h using a Leica SP8 AOBS confocal microscope with a 10x HC PL Fluotar lens (NA=0.3). Analysis of spheroid imaging data has been described before ^11^. Briefly, to obtain measurements of SIL density and spheroid dead volumes, raw data was pre-processed and semi-automatically analysed using a custom-written Cancer Segmentation workflow for the Fiji ^56^ plugin, MIA (v0.9.26) and its MIA_MATLAB (v1.1.1) package, available at GitHub via Zenodo: http://doi.org/10.5281/zenodo.2656513 and http://doi.org/10.5281/zenodo.4769615, respectively. The corresponding .mia workflow files are available at https://doi.org/10.5281/zenodo.5511888. The imaged stacks were mirrored and concatenated along the z-axis to produce pseudo-complete spheroids. These spheroids were binarised and segmented using connected-components labelling ^57^. Spheroids were fit with alpha shapes ^58^ using the MATLAB implementation. Adjacent spheroids were separated with a distance-based watershed transform ^57^. T-cells and dead volumes were individually segmented from their respective fluorescence channels using similar threshold and labelling-based processes, albeit without the alpha shape step.

As a control for minimal efficacy of CL4 killing, only experiments with an average increase in DRAQ7^+^ volume relative to 2h of 100,000 µm^3^ in the CL4 F-tractin-GFP condition were included. Comparing the ratio of CTL to accessible tumor cell surface area between the 2D and 3D spheroid killing assays, limiting numbers of CTL in tissue culture required the execution of the 2D killing assays at about a tenth of the ratio used in spheroids. Therefore, limiting activation conditions are reached sooner. This may apply to 1G4 TCR/TIM3-GFP and 1G4 TCR/F-tractin-GFP-transduced human CTL, where only 5% of the sorted CTL expressed detectable amounts of 1G4 TCR.

### Flow cytometry staining

Cells to be stained were resuspended in PBS at a concentration of 1×10^6^ cells/ml. 2.5×10^5^-1×10^6^ cells for each condition resuspended in 100μl PBS per tube with 1μl/100 μl Zombie Aqua Fixable Live Cell Detection reagent (BioLegend). Tubes were incubated for 15min in the dark at room temperature. Cells were washed in 3ml FACS buffer and resuspended in 100μl per tube FcBlock (BD Biosciences) for 15min at 4°C. Cells were washed in 3ml FACS buffer, pelleted and resuspended in 100μl FACS buffer per tube with antibody at the required concentration (antibody section above) and then incubated for 30min at 4°C. Antibody concentration was determined by titration using 5 concentrations centered around the manufacturer’s recommended protocol. Cells were washed in 3ml FACS buffer before being fixed in 1% paraformaldehyde and analyzed within 5 days using a Fortessa Flow Cytometer and BD FACSDiva Software (BD Biosciences).

For intracellular staining of Renca tumor cells for galectin9, cells were fixed and permeabilized using BD Biosciences ICCS Kit following the manufacturer’s protocol. Briefly, cells were resuspended in 100μl PBS per tube with 1μl/100μl Zombie Aqua Fixable Live Cell Detection reagent (BioLegend). Cells were fixed in 250 µl of Fixation/Permeabilization solution (BD Biosciences ICCS Kit) for 20min at 4°C. Cells were stained in 50µL of BD Perm/Wash™ buffer with 5µg/ml anti-mouse galectin9 mAb for 30min at 4°C. Cells were analyzed within 5 days using a Fortessa Flow Cytometer and BD FACSDiva Software (BD Biosciences). Flow cytometric data were analyzed using FlowJo™ (Treestar) software. Gating was performed using fluorescence minus one (FMO) samples for each antibody stain.

### Statistics and Reproducibility

The Power of the *in vivo* experiment was designed to reach >80%. Experimental group size was determined using the equation: *n = (2/(standardized difference2)) x cp,power,* where n = sample size per group determined using the formula, d = standardized difference = measurable difference in tumor volume / standard deviation, cp,power = constant for p<0.01 and power at 80% defined using standard Altman’s Nomogram = 11.7.

Samples were compared using independent sample t-tests for two sample comparisons. To determine the effect of one or more independent variables on one dependent variable across >2 groups, One-way and Two-way ANOVA were used once a normal distribution of the data has been confirmed. Where proportions were compared, a proportion’s z-test was used. SPSS statistics and Prism were used to execute analyses.

### Data and software availability

Image analysis workflow files are available at Zenodo as detailed in the methods section. Spheroid imaging data of 500Gb are available upon request.

