## Supplementary figures for "TIM3 is a context-dependent co-regulator of cytotoxic T cell function"

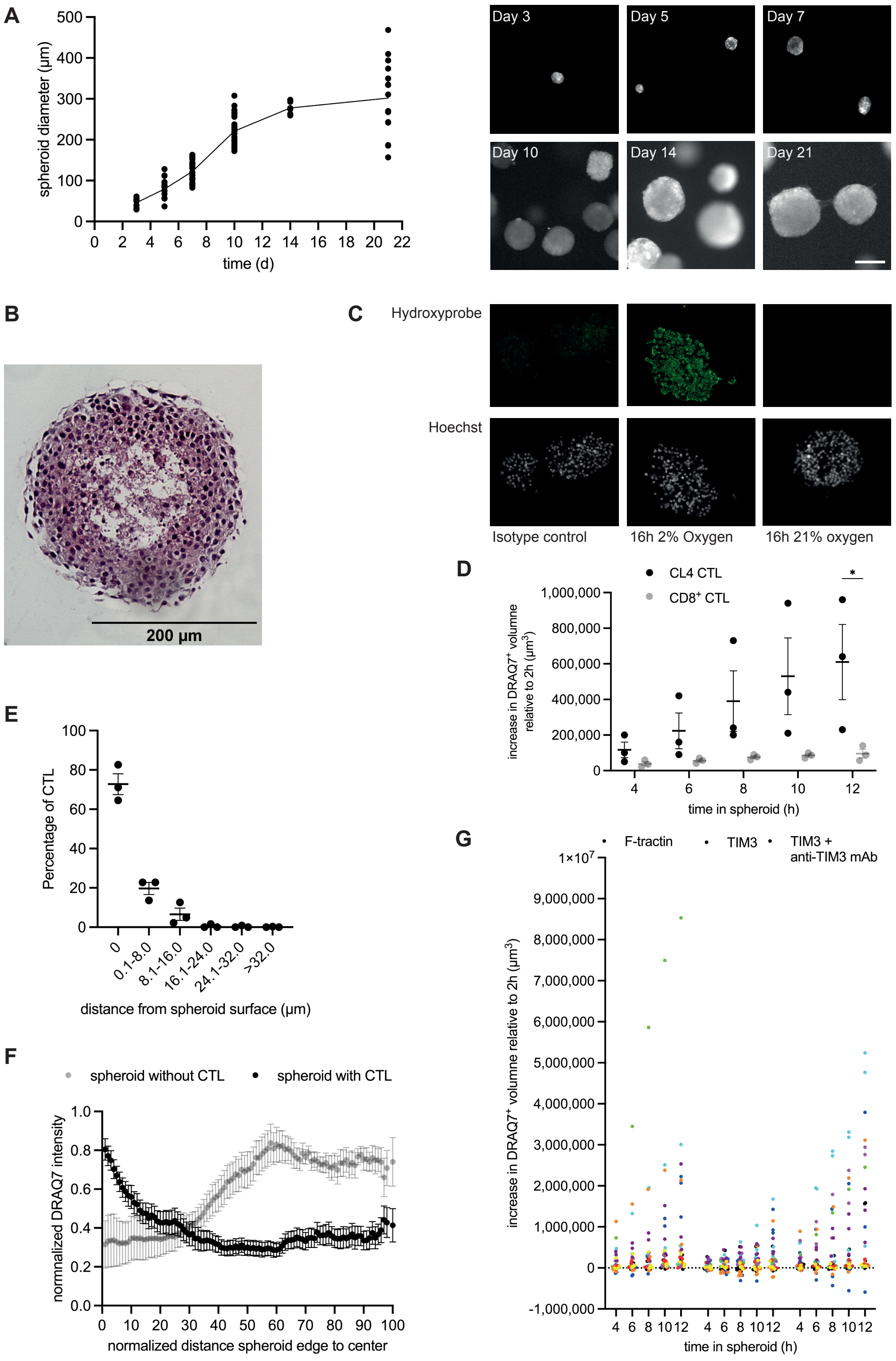
A
spheroid diameter (µm)
time (d)
Day 3
Day 5
Day 7
Day 10
Day 14
Day 21
B
200 µm
C
Hydroxyprobe
Hoechst
Isotype control
16h 2% Oxygen
16h 21% oxygen
D
CL4 CTL
CD8+ CTL
increase in DRAQ7+ volume relative to 2h (µm3)
time in spheroid (h)
E
Percentage of CTL
distance from spheroid surface (µm)
0
0.1-8.0
8.1-16.0
16.1-24.0
24.1-32.0
>32.0
F
spheroid without CTL
spheroid with CTL
normalized DRAQ7 intensity
normalized distance spheroid edge to center
G
F-tractin
TIM3
TIM3 + anti-TIM3 mAb
increase in DRAQ7+ volume relative to 2h (µm3)
time in spheroid (h)

**Figure S1 – Spheroid characterization**

**A** RencaHA tdTomato spheroid growth curve with representative images. N=3 independent experiments. **B** Representative midsection of a day 10 RencaHA tdTomato spheroid after fixation, paraffin embedding and H&E staining. **C** Representative hypoxia staining experiment. RencaHA tdTomato spheroids were incubated in hypoxic (2%$O_2$) or normoxic incubators for 16h and incubated with pimonidazole hydrochloride (HCl) for the last 2h before fixation, paraffin embedding and staining with Hydroxyprobe antibody conjugated to FITC. Cell nuclei were stained with Hoechst 33258. **D** CL4 CTL or BALB/c $CD8^+$ CTL expanded with anti-CD3/anti-CD28 beads ($CD8^+$ CLT) were cocultured with RencaHA tdTomato spheroids incubated with $K^d$HA peptide for 12h with images acquired every 2h. Spheroid death, as measured by the increase in $DRAQ7^+$ spheroid volume, is shown. Each data point is an independent experiment (N=3). **E,F** RencaHA tdTomato spheroids were incubated with or without CL4 CTL expressing F-tractin-GFP for 12h in the presence of DRAQ7. E Percentage of SIL at the given distance from the spheroid surface is given. N=3 independent experiments. F Relative DRAQ7 fluorescence intensity is given as a function of distance from the spheroid edge, normalized to the spheroid radius. 16, 6 spheroids from N=4, 2 independent experiments. **G** Single spheroid data for Fig. 1C. Spheroids from the same experiment share their color across all experimental conditions. * $p < 0.05$; p values calculated using two-way ANOVA

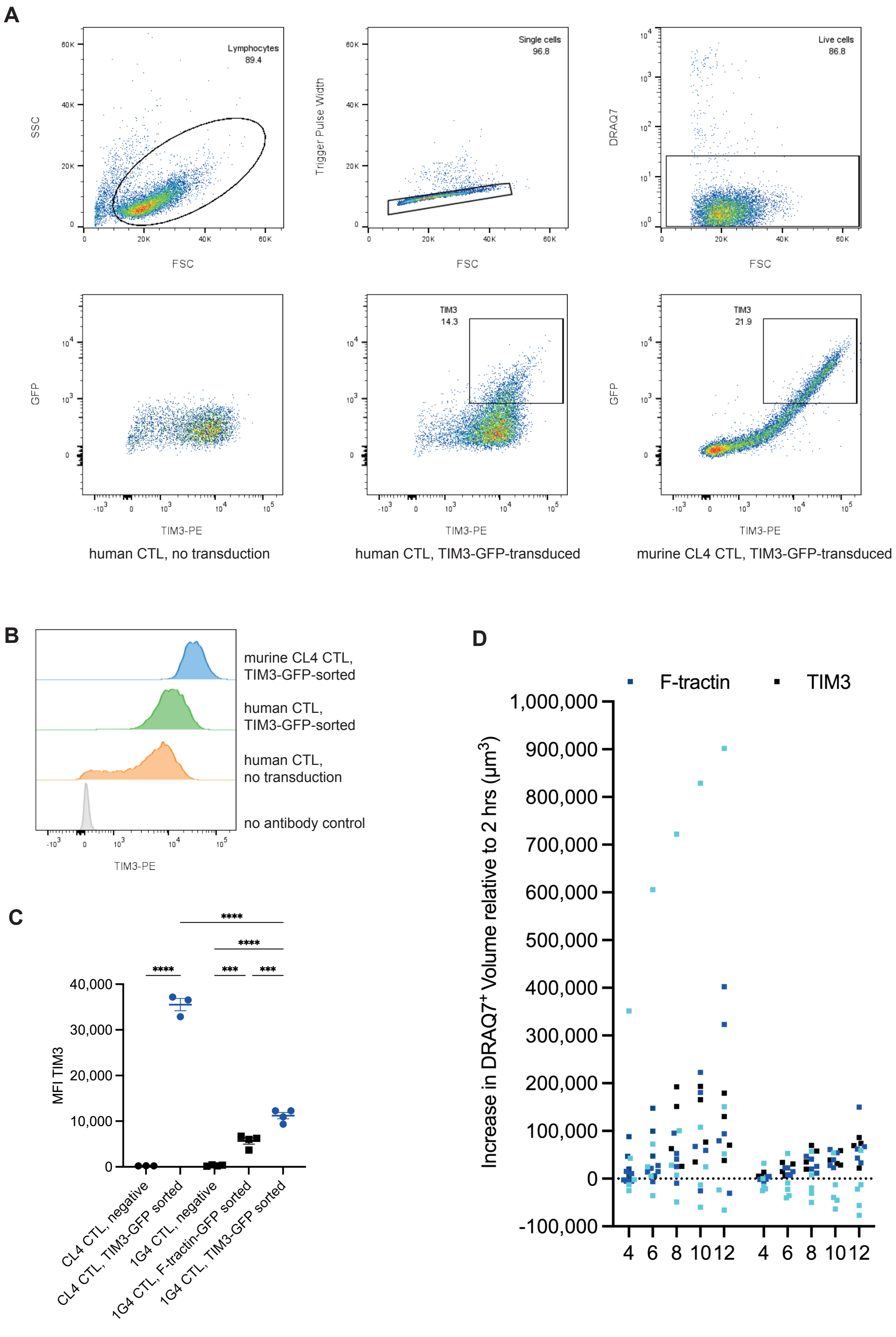
A
Lymphocytes
89.4
SSC
FSC
Single cells
96.8
Trigger Pulse Width
Live cells
86.8
DRAQ7
GFP
TIM3-PE
TIM3
14.3
TIM3
21.9
human CTL, no transduction
human CTL, TIM3-GFP-transduced
murine CL4 CTL, TIM3-GFP-transduced
B
murine CL4 CTL,
TIM3-GFP-sorted
human CTL,
TIM3-GFP-sorted
human CTL,
no transduction
no antibody control
TIM3-PE
C
MFI TIM3
40,000
30,000
20,000
10,000
0
****
****
****
***
***
CL4 CTL, negative
CL4 CTL, TIM3-GFP sorted
1G4 CTL, negative
1G4 CTL, F-tractin-GFP sorted
1G4 CTL, TIM3-GFP sorted
D
F-tractin
TIM3
Increase in DRAQ7+ Volume relative to 2 hrs (µm3)
1,000,000
900,000
800,000
700,000
600,000
500,000
400,000
300,000
200,000
100,000
0
-100,000
4
6
8
10
12
4
6
8
10
12

**Figure S2 – TIM3 expression and human single spheroid data**

**A** Top row: gating strategy for the flow cytometry analysis of T cell cultures. Bottom row: Representative anti-TIM3 flow cytometry data of day 8 human CTL without lentiviral transduction (left), human CTL transduced to express the 1G4 TCR with TIM3-GFP (middle), murine CL4 CTL transduced to express TIM3-GFP, with TIM3-GFP sort gates. 3-4 independent experiments. **B** Representative flow cytometry histograms of TIM3 staining of day 8 human CTL without lentiviral transduction, human CTL sorted for TIM3-GFP expression as gated in A, mouse CL4 CTL sorted for TIM3-GFP as gated in A. 3-4 independent experiments. **C** MFI of anti-TIM3 staining of CL4 CTL sorted for TIM3-GFP as in A and the GFP-negative cells from the same population, of 1G4 CTL sorted for TIM3-GFP as in A, sorted for F-tractin-GFP through the same gate and the negative cells from the same population. 3-4 independent experiments. **D** Single spheroid data for Fig. 2B. Spheroids from the same experiment share their color across all experimental conditions. *** $p<0.001$, **** $p < 0.0001$; p values calculated using one-way ANOVA

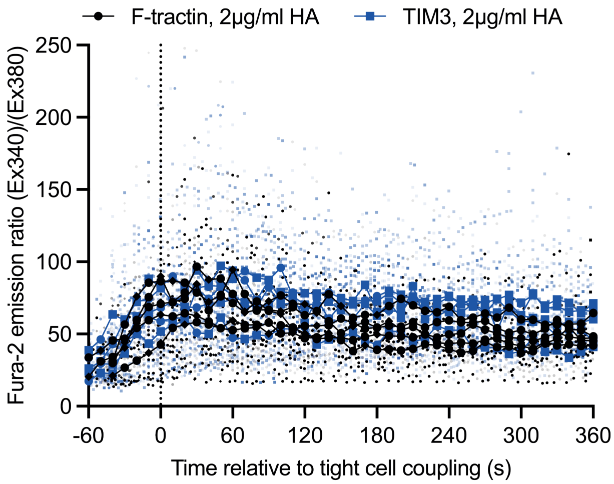


**Figure S3 – TIM3 overexpression does not alter CL4 CTL calcium signaling**
Single cell and individual experiment average data are given for Fig. 3F. Same experiments share the same symbol.

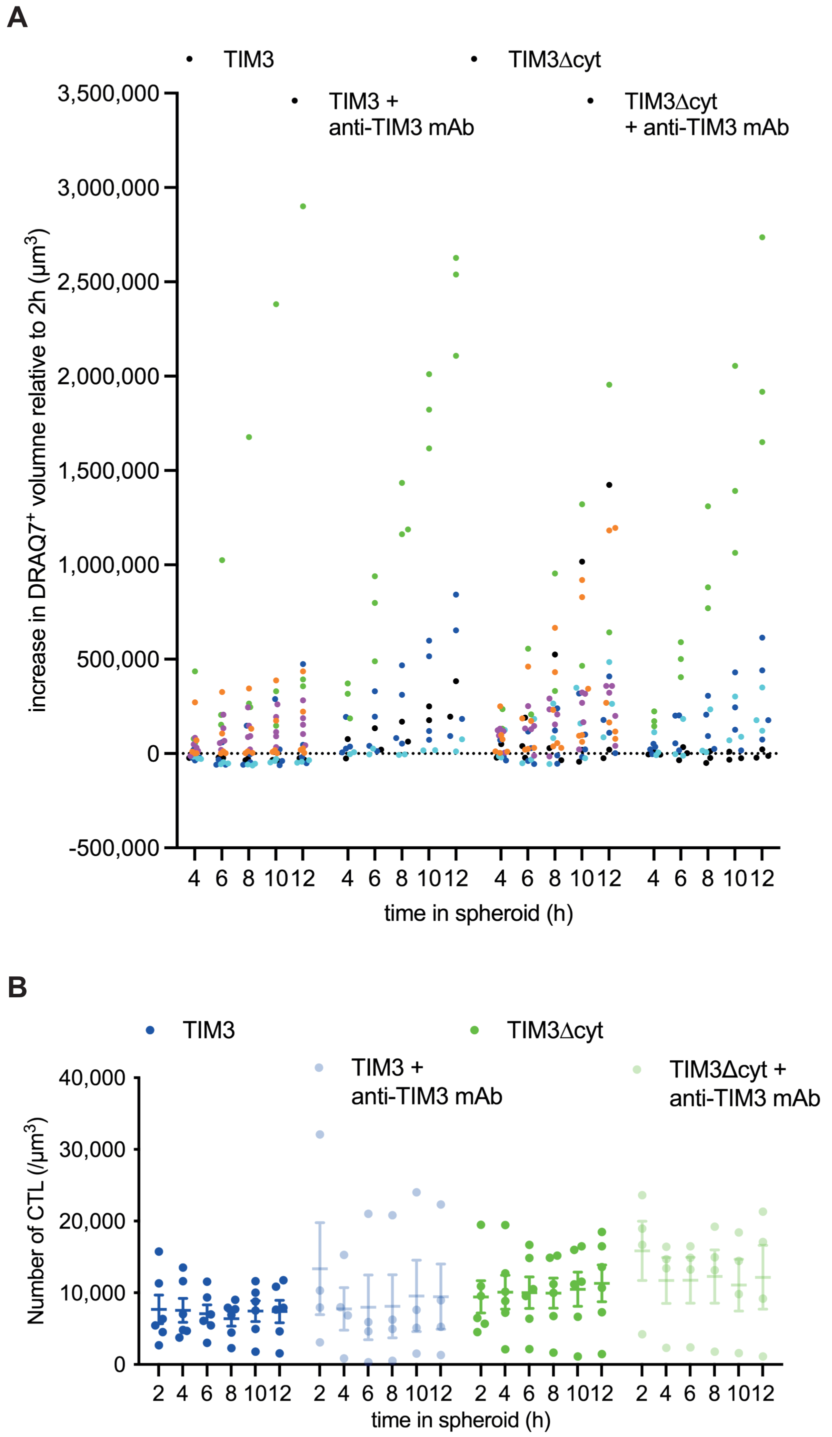


**Figure S4 – TIM3 co-regulatory function is only partially dependent on its cytoplasmic domain**
**A** Single spheroid data for Fig. 4A. Spheroids from the same experiment share their color across all experimental conditions. **B** SIL densities are shown with the mean ± SEM for the same experiments as in Fig. 4A. No significant differences.

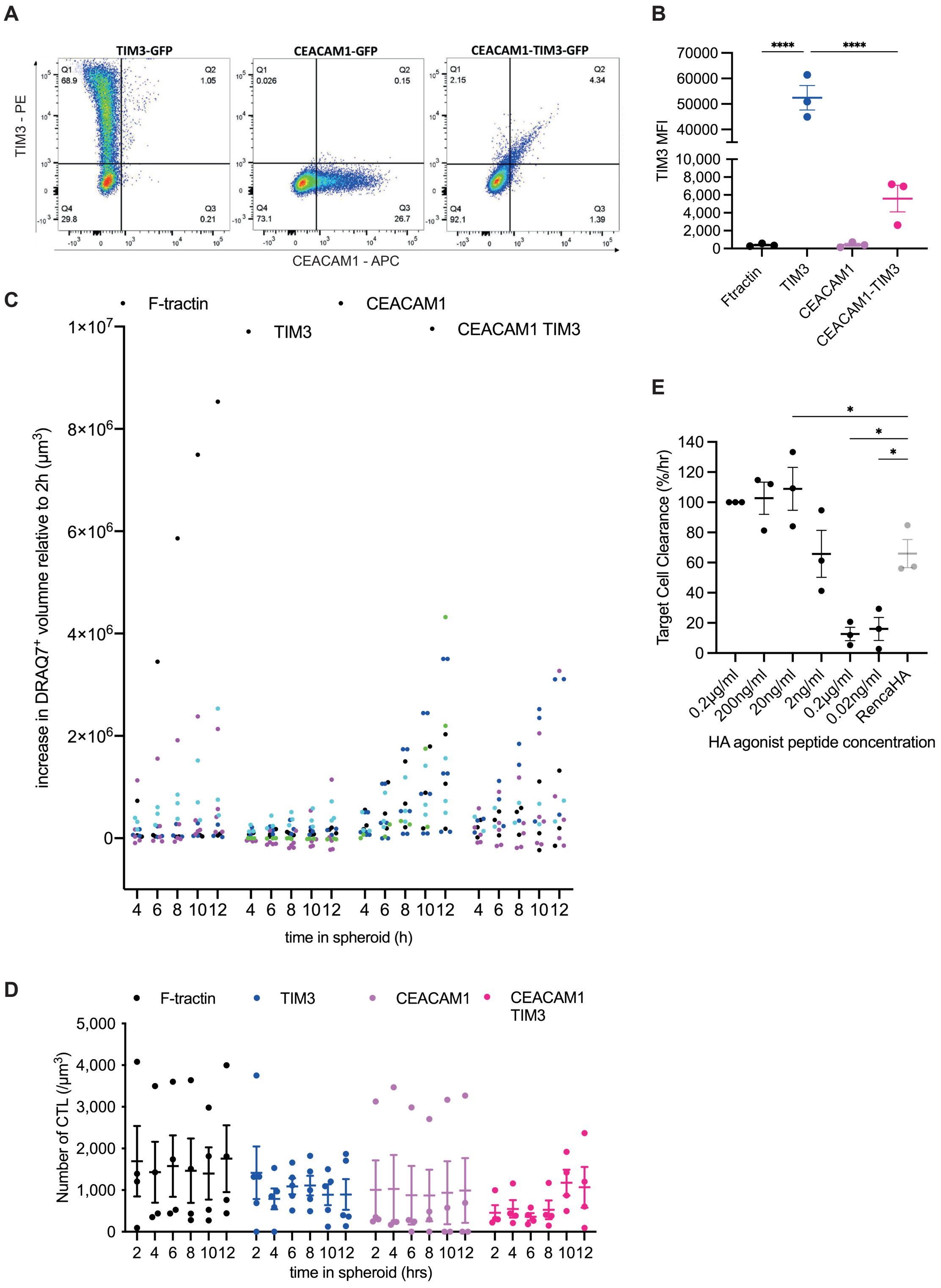
A
TIM3-GFP
CEACAM1-GFP
CEACAM1-TIM3-GFP
TIM3 - PE
CEACAM1 - APC
Q1 68.9
Q2 1.05
Q4 29.8
Q3 0.21
Q1 0.026
Q2 0.15
Q4 73.1
Q3 26.7
Q1 2.15
Q2 4.34
Q4 92.1
Q3 1.39
B
TIM3 MFI
****
****
Ftractin
TIM3
CEACAM1
CEACAM1-TIM3
C
F-tractin
TIM3
CEACAM1
CEACAM1 TIM3
increase in DRAQ7⁺ volumne relative to 2h (μm³)
time in spheroid (h)
D
F-tractin
TIM3
CEACAM1
CEACAM1 TIM3
Number of CTL (/μm³)
time in spheroid (hrs)
E
Target Cell Clearance (%/hr)
0.2μg/ml
200ng/ml
20ng/ml
2ng/ml
0.2μg/ml
0.02ng/ml
RencaHA
HA agonist peptide concentration

**Figure S5 – CEACAM1 expression in cis inhibits TIM3 function**

**A** Representative flow cytometry data of CL4 CTL expressing TIM3-GFP, CEACAM1-GFP or TIM3-GFP together with CEACAM1 stained for TIM3 and CEACAM1. **B** MFI of TIM3 expression in CL4 CTL transduced to express F-tractin-GFP, TIM3-GFP, CEACAM1-GFP or TIM3-GFP together with CEACAM1 sorted for GFP expression of at least ten-fold over negative, as used in the functional experiments. **C** Single spheroid data for Fig. 5B. Spheroids from the same experiment share their color across all experimental conditions. **D** SIL densities are shown with the mean ± SEM for the same experiments as in Fig. 5B. No significant differences. **E** *In vitro* killing of Renca cells (black) incubated with the indicated concentrations of $K^d$HA peptide as compared to RencaHA cells (grey) not incubated with exogenous peptide by CL4 CTL. Average cell death rate calculated as percentage decrease in area covered by tumor cells per hour are means ± SEM from 3 independent experiments. * $p < 0.05$, **** $p<0.0001$, p values calculated using one-way ANOVA (in E only shown for the comparison between RencaHA cells and the lowest four peptide concentrations).

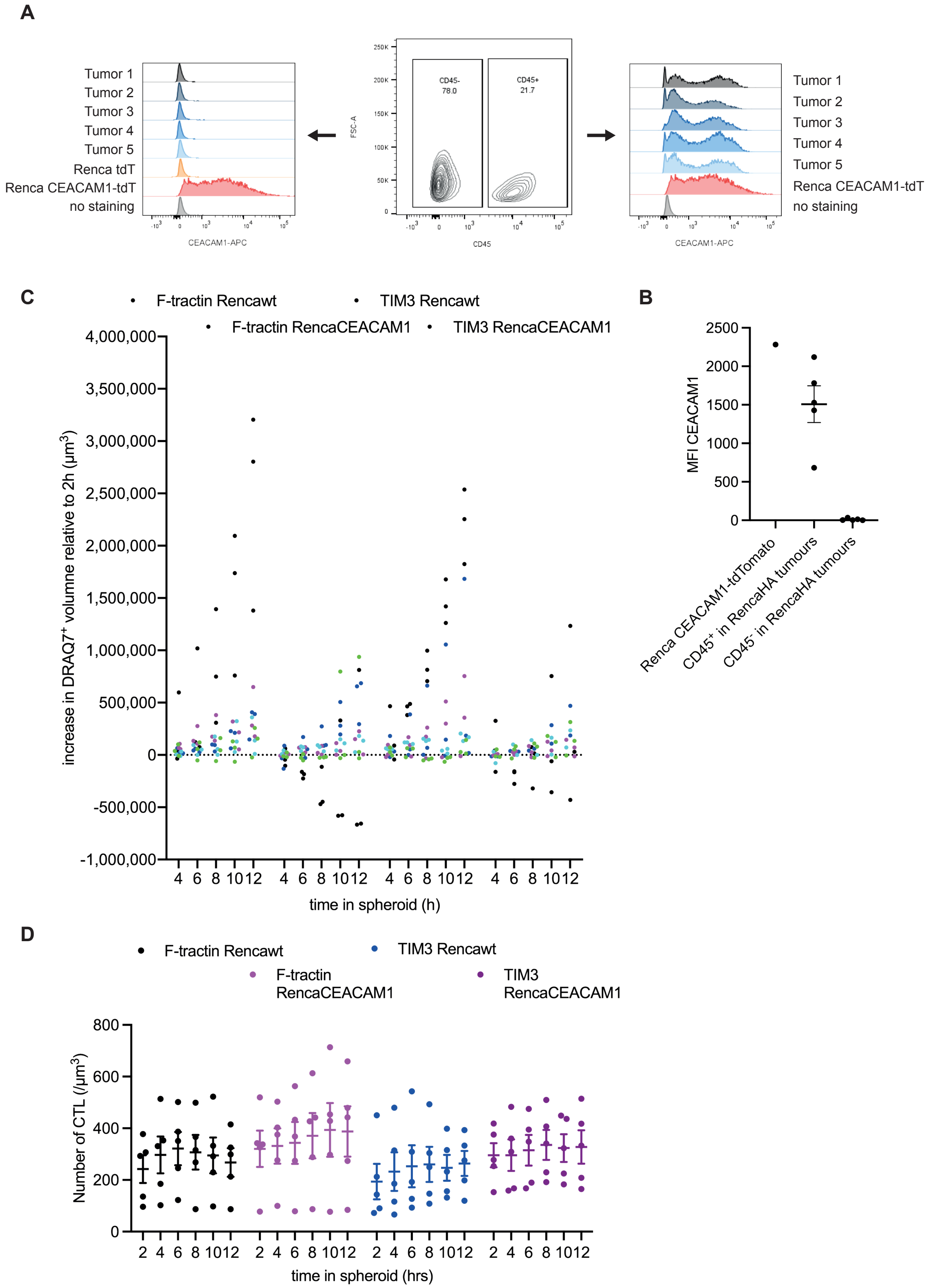
A
Tumor 1
Tumor 2
Tumor 3
Tumor 4
Tumor 5
Renca tdT
Renca CEACAM1-tdT
no staining
CEACAM1-APC
CD45-
78.0
CD45+
21.7
FSC-A
CD45
Tumor 1
Tumor 2
Tumor 3
Tumor 4
Tumor 5
Renca CEACAM1-tdT
no staining
CEACAM1-APC
C
F-tractin Rencawt
TIM3 Rencawt
F-tractin RencaCEACAM1
TIM3 RencaCEACAM1
increase in DRAQ7+ volume relative to 2h (µm3)
time in spheroid (h)
B
MFI CEACAM1
Renca CEACAM1-tdTomato
CD45+ in RencaHA tumours
CD45- in RencaHA tumours
D
F-tractin Rencawt
TIM3 Rencawt
F-tractin RencaCEACAM1
TIM3 RencaCEACAM1
Number of CTL (/µm3)
time in spheroid (hrs)

**Figure S6 – CEACAM1 in trans is an inhibitory TIM3 ligand**

**A** Flow cytometry data of cell surface staining for CEACAM1 of Renca cells expressing CEACMA1-tdTomato, Renca cells isolated from subcutaneous tumors as $CD45^{-}$ cells and immune cells isolated from subcutaneous tumors as $CD45^{+}$ cells. **B** MFI ± SEM of CEACAM1 expression from A. N=5 tumors. **C** Single spheroid data for Fig. 6C. Spheroids from the same experiment share their color across all experimental conditions. **D** SIL densities are shown with the mean ± SEM for the same experiments as in Fig. 6C. No significant differences.

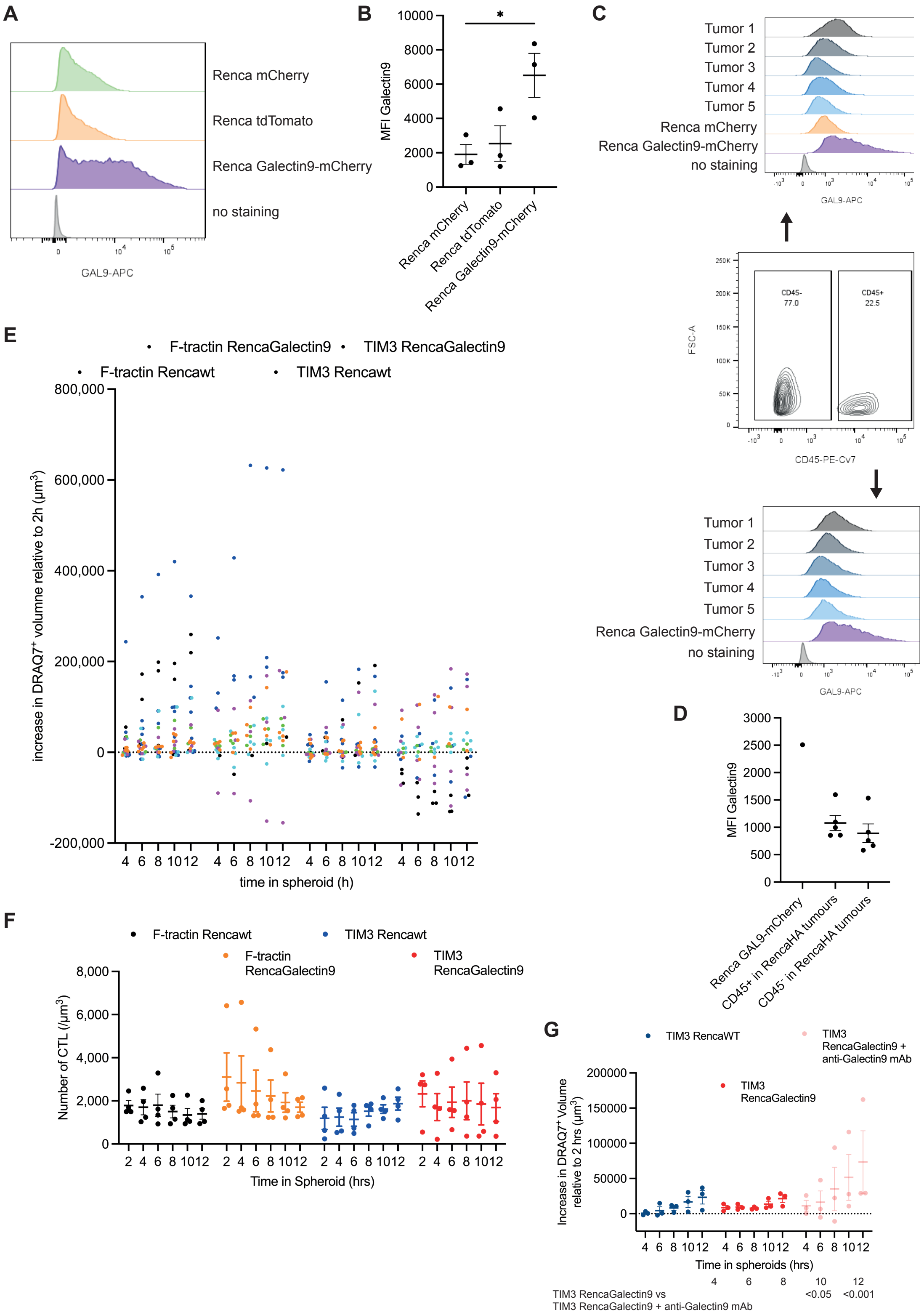

A
Renca mCherry
Renca tdTomato
Renca Galectin9-mCherry
no staining
GAL9-APC
B
MFI Galectin9
Renca mCherry
Renca tdTomato
Renca Galectin9-mCherry
C
Tumor 1
Tumor 2
Tumor 3
Tumor 4
Tumor 5
Renca mCherry
Renca Galectin9-mCherry
no staining
GAL9-APC
CD45-
77.0
CD45+
22.5
FSC-A
CD45-PE-Cv7
Tumor 1
Tumor 2
Tumor 3
Tumor 4
Tumor 5
Renca Galectin9-mCherry
no staining
GAL9-APC
D
MFI Galectin9
Renca GAL9-mCherry
CD45+ in RencaHA tumours
CD45- in RencaHA tumours
E
F-tractin RencaGalectin9
TIM3 RencaGalectin9
F-tractin Rencawt
TIM3 Rencawt
increase in DRAQ7+ volumne relative to 2h (µm3)
time in spheroid (h)
F
F-tractin Rencawt
TIM3 Rencawt
F-tractin RencaGalectin9
TIM3 RencaGalectin9
Number of CTL (/µm3)
Time in Spheroid (hrs)
G
TIM3 RencaWT
TIM3 RencaGalectin9 + anti-Galectin9 mAb
TIM3 RencaGalectin9
Increase in DRAQ7+ Volume relative to 2 hrs (µm3)
Time in spheroids (hrs)
TIM3 RencaGalectin9 vs
TIM3 RencaGalectin9 + anti-Galectin9 mAb
4
6
8
10 <0.05
12 <0.001

**Figure S7 – Galectin9 is an inhibitory TIM3 ligand**

**A** Representative flow cytometry data of Renca cells and transfectants thereof as indicated stained for Galectin9 after fixation and permeabilization. **B** MFI ± SEM of Galectin9 staining of Renca cells and transfectants thereof as indicated after fixation and permeabilization. N=3 independent experiments. **C** Flow cytometry data of Galectin9 staining after fixation and permeabilization of Renca cells expressing Galectin9-mCherry, Renca cells isolated from subcutaneous tumors as $CD45^{-}$ cells and immune cells isolated from subcutaneous tumors as $CD45^{+}$ cells. **D** MFI ± SEM of Galectin9 expression from C. N=5 tumors. **E** Single spheroid data for Fig. 7B. Spheroids from the same experiment share their color across all experimental conditions. **F** SIL densities are shown with the mean ± SEM for the same experiments as in Fig. 7B. No significant differences. **G** CL4 CTL retrovirally transduced to express TIM3-GFP cocultured with RencaHA tdTomato or RencaHA Galectin9-tdTomato spheroids ± 10µg/ml anti-galectin9 mAb incubated with $K^{d}$HA peptide for 12h with images acquired every 2h. Each data point is an independent experiment (N=3) with a total of 10 spheroids analyzed per condition. Spheroid death, as measured by the increase in $DRAQ7^{+}$ spheroid volume, is shown. Significance of differences between conditions at indicated time points is given in the table below. * $p<0.05$ , p values calculated using one-way (B) or two-way (G) ANOVA

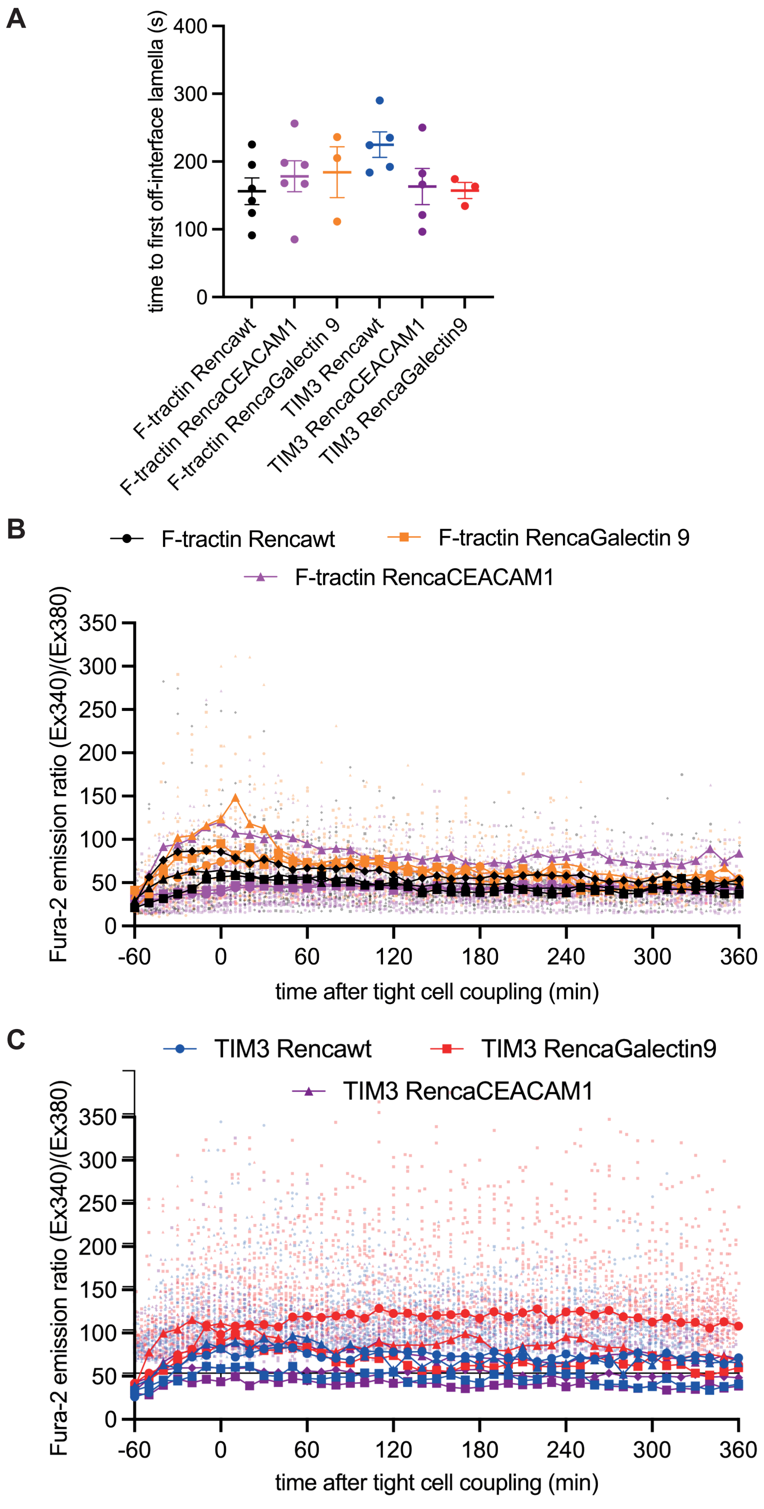


**Figure S8 – CTL morphology relates to cytolysis and calcium signaling to IFNγ secretion**
**A** For the same cell couples as analyzed in Fig. 8A-C, time of first off-synapse lamella. **B, C** Single cell and individual experiment average data are given for Fig. 8I, J. Same experiments share the same symbol across all conditions.

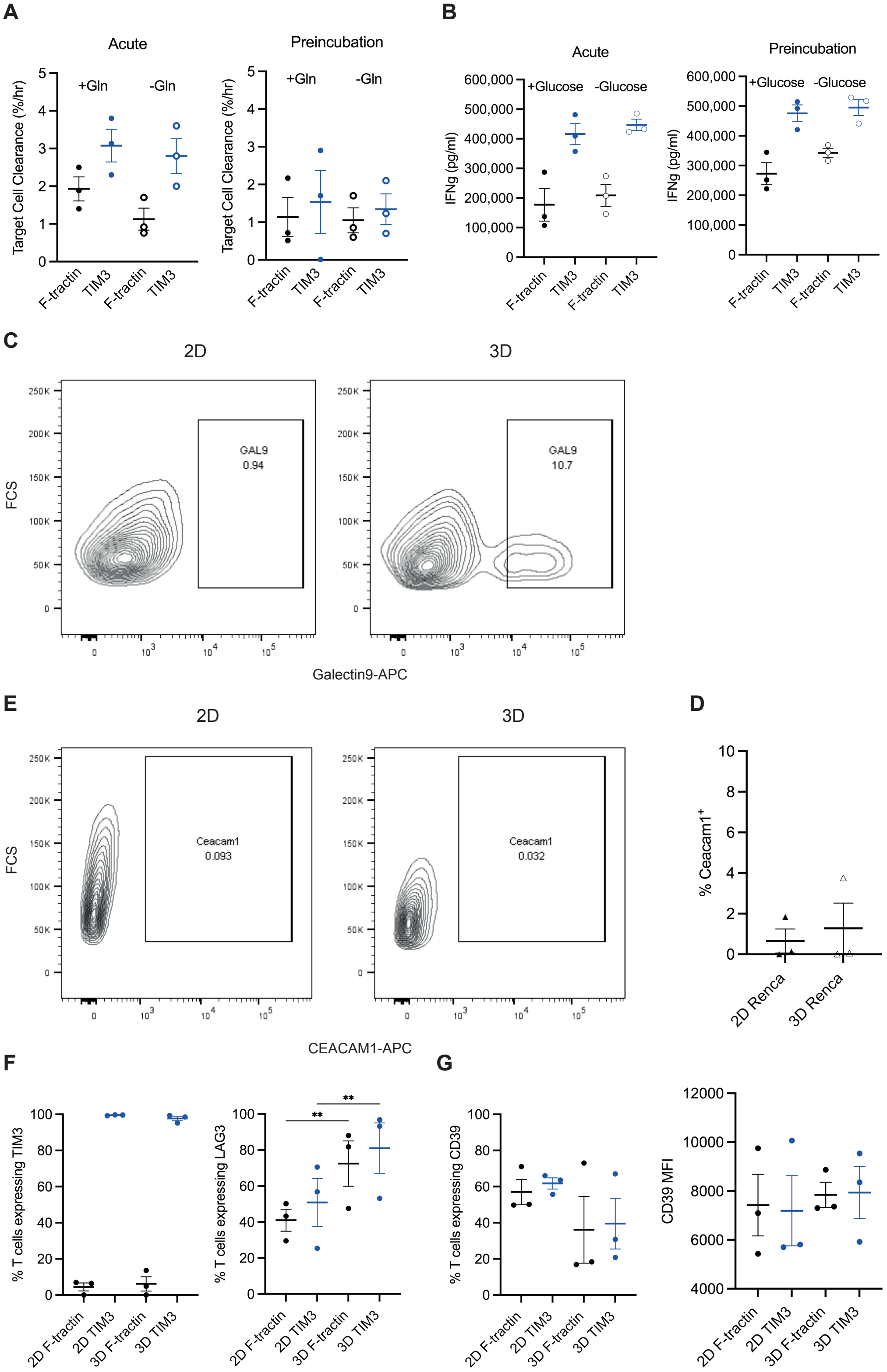

A
Acute
Preincubation
Target Cell Clearance (%/hr)
+Gln
-Gln
F-tractin
TIM3
B
IFNg (pg/ml)
+Glucose
-Glucose
C
2D
3D
FCS
GAL9
0.94
10.7
Galectin9-APC
E
Ceacam1
0.093
0.032
CEACAM1-APC
D
% Ceacam1+
2D Renca
3D Renca
F
% T cells expressing TIM3
% T cells expressing LAG3
2D F-tractin
2D TIM3
3D F-tractin
3D TIM3
G
% T cells expressing CD39
CD39 MFI

**Fig. S9 – The physiological environment for TIM3 function is more inhibitory in spheroids**

**A** Killings assays for Fig. 9B. **B** IFN$\gamma$ in killing assay supernatants for Fig. 9C. **C** Representative RencaHA Galectin9 staining data for Fig. 9F. **D** Percentage RencaHA cells with detectable CEACAM1 after overnight culture in 2D or spheroids ('3D'). 3 independent experiments. **E** Representative RencaHA CEACAM1 staining data for D. **F** Percent CL4 T cells positive for TIM3 or LAG3 from the experiments in Fig. 9G, I. **G** Percent CL4 T cells positive for CD39 and mean fluorescence intensity (MFI) of staining for CD39 of CL4 CTL expressing F-tractin-GFP or TIM3-GFP after overnight culture with 3D spheroids ('3D') or 2D culture with IL-2 ('2D'). 3 independent experiments. ** $p < 0.01$, p values calculated using one-way ANOVA
